# Germinal centers can persist for up to one year with declining somatic hypermutation and ongoing plasma cell output

**DOI:** 10.64898/2026.10.13.758939

**Authors:** Henry J. Sutton, Ester Marina-Zárate, Tasha K. Altheide, Arpan Pradhan, Amit A. Upadhyay, Iszac Burton, Michael Bick, Agnes Walsh, Mariane B. Melo, Jenny Tran, Nicole Phelps, Danny Lu, Diana Goodwin, Wen-Hsin Lee, Kipchoge N Randall, Andy S. Tran, Kesha Hyacinth, Brandon S. Healy, Deuk Lim, Diane G. Carnathan, Christopher A. Cottrell, Lars Hangartner, Dennis R. Burton, Guido Silvestri, Andrew B. Ward, Gabriel Ozorowski, Steven E. Bosinger, Sudhir P. Kasturi, William R. Schief, Darrell J. Irvine, Shane Crotty

## Abstract

The total duration of primary germinal center (GC) responses, and how B cell evolution changes as these responses mature, remain poorly defined. Here, longitudinal sampling of rhesus macaques immunized with HIV Envelope protein revealed that a single slow-delivery prime sustained ongoing GCs for nearly a year without additional antigen. Somatic hypermutation (SHM) accumulated throughout these prolonged responses, but its rate declined substantially over time, independent of both priming strategy and boosting. Over the same period, subdominant non-base Env specificities became increasingly represented within the GC response. Delaying boosting until after primary GC contraction increased recruitment of recalled memory B cells into the GC response. Bone marrow plasma cells were generated throughout prolonged GC maturation, with the timing of their generation modulated by booster timing. Together, these findings reveal how GCs evolve across nearly a year and how this prolonged maturation shapes the memory and plasma cell compartments that underpin long-term humoral immunity.

## INTRODUCTION

Germinal centers (GCs) are dynamic sites of B cell evolution that progressively shape the affinity, specificity, and diversity of antibody responses. This evolution is driven by iterative cycles of somatic hypermutation (SHM), proliferation, and affinity-dependent selection^1–4^. Murine studies led to the longstanding view that vaccine-induced GC responses typically resolve within weeks^5–7^, with longer persistence observed in some infection settings^8,9^. However, accumulating evidence in human and non-human primates (NHPs) indicates that GCs can persist considerably longer following vaccination or infection^10–18^. Much of this insight has come from the use of longitudinal lymph node (LN) fine needle aspiration (FNA), which enables repeated sampling of the same GC responses over extended periods^19^, identifying GCs that persist for at least six months following vaccination against SARS-CoV-2 and seasonal influenza in humans^10,12–14^. Comparable persistence has been achieved in rhesus macaques (RMs) through slow-delivery immunization, in which antigen is administered over several days rather than as a single bolus^20,21^. A single escalating-dose (ED) prime with HIV Envelope (Env) protein sustained Env⁺ GCs for six months without further antigen administration^16^. These GCs remained functionally active, with B cell lineages maintained across the six-month response acquiring up to ∼1,000-fold improvements in binding affinity. Notably, in each of these studies, antigen-specific GC B cells (B_GC_ cells) remained detectable when sampling ended, leaving the true lifespan of vaccine-induced GCs undefined.

Longer-lived GCs potentially provide greater opportunity for B cell clones to accumulate SHM, but whether the rate of SHM remains stable over the lifespan of a GC, particularly during prolonged responses, remains unknown. Prolonged GC activity may also alter the specificity of the response. For complex antigens, antibody-mediated masking of immunodominant epitopes can alter competition within the GC and favor B cells recognizing subdominant epitopes, including those that may be important targets for protective or broadly reactive antibodies^22–24^. Longer-lived GCs, by providing more opportunity for these competitive processes to operate, might be expected to diversify the responding repertoire across a broader range of epitopes.

The persistence of GC activity for many months also has important implications for vaccination strategies involving repeated immunization. This is especially relevant to germline-targeting sequential-immunization HIV vaccine regimens, which use a succession of immunogens to guide broadly neutralizing antibody (bnAb) precursor B cells through intermediaries to recognition of diverse HIV strains^25,26^. Because boosts in these regimens are typically given while the GC response to the preceding immunogen is still active, a booster may be delivered in the presence of both an active primary GC and an evolving memory B cell (B_Mem_) compartment. Whether boosting primarily refuels cells already participating in the ongoing GC^27^ or recalls B_Mem_ cells into de novo secondary GCs, and how this balance varies with booster timing, remains unclear.

The success of sequential vaccination ultimately depends on whether progressively matured B cells are recruited into the long-lived plasma cell (B_PC_) compartment, where they can sustain protective bnAb titers. However, when long-lived B_PCs_ are generated over the course of an immune response remains ill defined, particularly in humans and NHPs, where vaccine-induced B_PC_ responses are difficult to track longitudinally. Studies in mouse models have suggested two potential patterns of B_PC_ generation during the primary response: B_PCs_ may be generated continuously throughout GC maturation^28–31^, or B_PC_ differentiation/accumulation may instead be concentrated toward later stages of the response^32,33^. An additional question is how antigen re-exposure alters this process. In sequential vaccination, where repeated boosts are required to progressively mature bnAb precursor lineages, the timing of boosting could influence which stages of the evolving B cell response are captured within the long-lived B_PC_ compartment. Defining how B_PC_ generation is regulated by both prolonged immune responses and antigen re-exposure may help inform the optimal timing of sequential immunizations.

Resolving how these processes unfold over time requires following both prolonged GC responses and their downstream B cell outputs across extended periods. Here, repeated sampling of LNs, blood, and bone marrow (BM) was performed in RMs receiving bolus or slow-delivery HIV Env priming. Booster immunizations were administered at different stages of these prolonged GC responses to examine the consequences of antigen re-exposure during GC maturation. Longitudinal analysis over one year established the duration and evolution of vaccine-induced GCs and determined how GC maturation and the timing of antigen re-exposure influence B_Mem_ recall and the BM B_PC_ compartment.

## RESULTS

### Study design

To define how priming strategy and booster timing shape prolonged GC responses and their downstream humoral output, we compared four immunization regimens in 32 Indian RMs (n = 8 per group) **(Fig. 1A)**. All animals received 100 μg of MD39 Env trimer formulated with 750 μg of saponin/MPLA nanoparticle (SMNP)^34^, administered bilaterally into the left and right mid-thighs. Groups 1 and 2 received a bolus prime, whereas Groups 3 and 4 were primed with the previously described 12-day slow-delivery escalating dose (ED)^16,21,35–38^. Group 1 was boosted at weeks 10 and 24, whereas Groups 2 and 3 were boosted at week 24 and Group 4 at week 52. Blood and serum, bilateral draining LN FNAs, and bone marrow (BM) aspirates were collected longitudinally to track Env-specific GC B cells (B_GC_), B_Mem_, B_PCs_, and antibody responses **(Fig. 1A)**.

**Figure 1.**
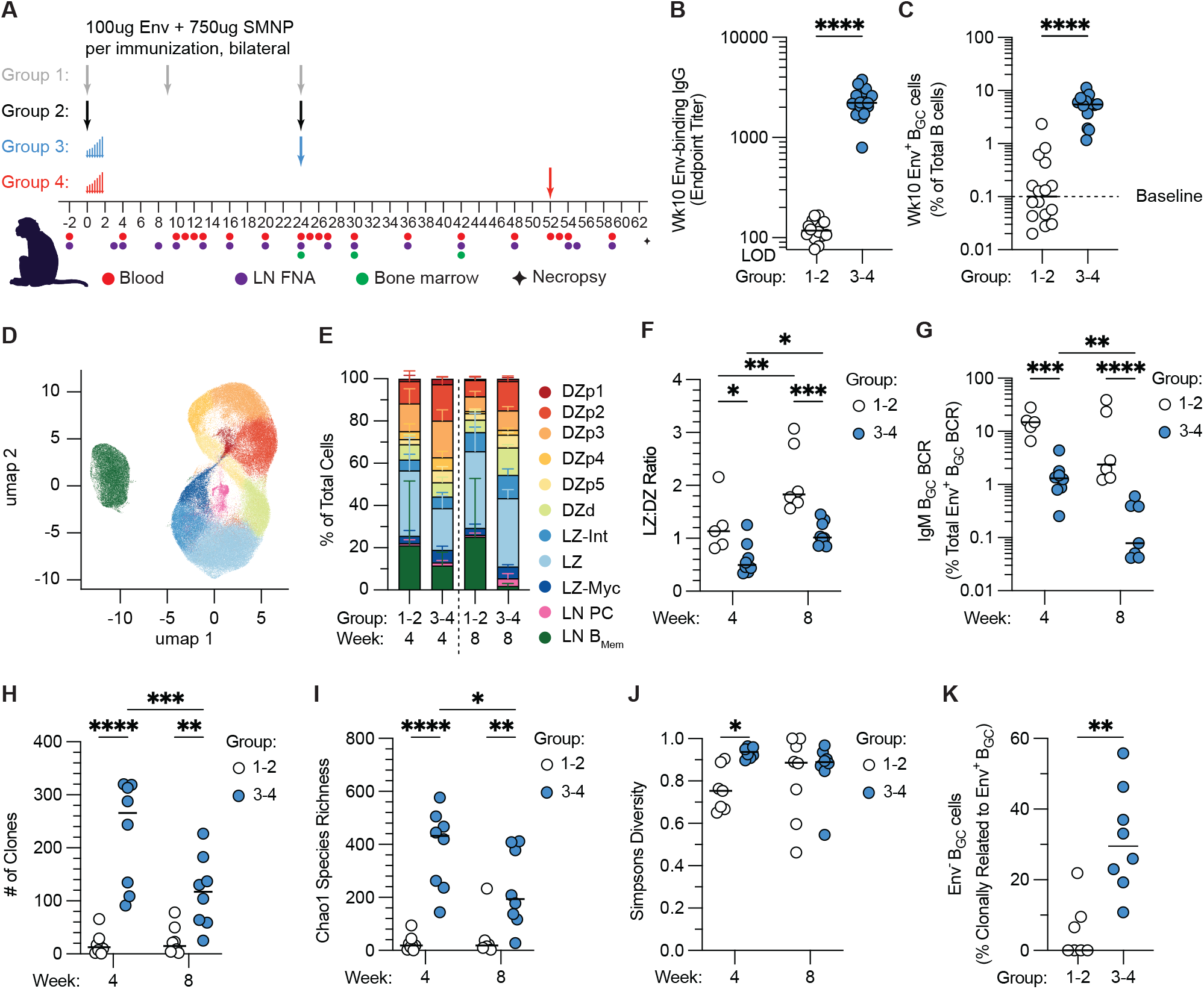
Slow-delivery priming generates larger and more diverse early germinal center responses. **(A)** Schematic of experimental design. Rhesus macaques received a total dose of 100 µg MD39 Env trimer formulated with 750 µg SMNP adjuvant split between the left and right thigh per immunization. Single arrows indicate bolus immunizations, whereas a series of increasing arrows indicates escalating-dose (ED) immunization. Blood, lymph node fine-needle aspirate (LN FNA), and bone marrow samples were collected at the indicated time points. **(B)** Env-specific serum IgG endpoint titers at week 10 following bolus (Groups 1 and 2) or ED (Groups 3 and 4) priming. The y-axis begins at the limit of detection (LOD). **(C)** Frequency of Env⁺ B_GC_ cells as a percentage of total B cells in LN FNAs at week 10. Dotted line indicates frequency at baseline (week -2). **(D)** uniform manifold approximation and projection (UMAP) of LN B cell clusters identified by single-cell RNA sequencing of LN FNAs. **(E)** Relative abundance of LN B cell clusters identified in **(D)** following bolus or ED priming at weeks 4 and 8. **(F)** Light zone-to-dark zone (LZ:DZ) ratio of Env⁺ B_GC_ cells at weeks 4 and 8. **(G)** Frequency of Env⁺ B_GC_ cells with IgM B cell receptors (BCRs) at weeks 4 and 8. **(H)** Total number of Env⁺ B_GC_ cell clones identified per animal at weeks 4 and 8. **(I)** Chao1 species richness of Env⁺ B_GC_ cell clonal repertoires at weeks 4 and 8. **(J)** Gini–Simpson diversity index of Env⁺ B_GC_ cell clonal repertoires at weeks 4 and 8. **(K)** Frequency of Env⁻ B_GC_ cells clonally related to Env⁺ B_GC_ cells at weeks 4 and 8. Statistical comparisons were performed using two-tailed Mann–Whitney tests for **(B)**, **(C)**, and **(K)**, and two-way ANOVA with multiple-comparisons correction for **(F–J)**. For analyses of B_GC_ phenotype and BCR characteristics in **(E– G),** samples with fewer than 30 Env⁺ B_GC_ sequences were excluded because of insufficient sampling depth. For **(I)**, samples for which Chao1 richness could not be reliably estimated were excluded. *P* values are indicated as *P* < 0.05 (\**), P < 0.01 (**), P < 0.001 (\*\*\**), and *P* < 0.0001 (****); non-significant comparisons are not shown. Horizontal bars indicate median.

### Slow-delivery priming elicits larger, more diverse germinal-center responses than bolus immunization

We first asked how antigen delivery kinetics influenced the magnitude and composition of the early GC response. At 10 weeks post prime, ED-primed RMs had significantly higher Env-binding IgG titers than bolus-primed RMs (*P* < 0.0001; **Fig. 1B**). Consistent with this, slow-delivery priming generated significantly larger Env^+^ GC responses in the draining LN than a bolus prime (median 5.47% vs. 0.10% of B cells; *P* < 0.0001; **Fig. 1C; Fig. S1A**), whereas Env⁺ GC frequencies in most bolus-primed RMs had returned to baseline levels by week 10.

To characterize these responses in greater depth, longitudinal single-cell RNA sequencing (scRNA-seq) with paired VDJ sequencing was performed on Env^+^ B cells from a representative subset of four animals per group (16 animals total). CD20⁺IgD⁻Env^+^ B cells were sorted from the left draining LN (LLN) at weeks 4, 8, 10, 16, 24, 48, 55 and 59, together with matched samples for the right draining LN (RLN) at week 4 (**Fig. S1B**). In addition, an Env⁻ population was sorted from the RLN at week 4 to identify low-affinity Env-specific clones. Following quality control, 175,490 LN B cells were retained for downstream transcriptomic analysis. Unsupervised clustering resolved 11 LN B cell populations comprising five proliferating dark zone (DZ) clusters (DZp1-5), one differentiating dark zone cluster (DZd)^39^, three light zone (LZ) populations consisting of a classical LZ cluster (LZ), a MYC-expressing LZ cluster (LZ-Myc), and an intermediate cluster between the LZ and DZ (LZ-Int), together with an LN plasma cell (LN B_PC_) cluster and an LN memory B cell (LN B_Mem_) cluster (**Fig. 1D, Fig. S1C-D**). This transcriptional classification enabled B_GC_ cells to be distinguished from LN B_Mem_ and LN B_PCs_ for all subsequent analyses. During the early response (weeks 4 and 8), ED-primed RMs contained a greater proportion of DZ cells than bolus-primed animals (**Fig. 1E**) resulting in a significantly lower LZ:DZ ratio following a slow-delivery prime at both weeks 4 (*P* = 0.03) and 8 (*P* = 0.0002; **Fig. 1F**). Although the ratio increased over time in both groups as the GC matured, ED-primed RMs maintained a proportionally larger DZ compartment, accompanied by a higher frequency of proliferating B_GC_ cells at both time points (**Fig. S1E-F**).

Paired VDJ sequencing performed alongside scRNA-seq enabled analysis of isotype usage within the GC response. Notably, bolus-primed RMs contained substantially more unswitched IgM^+^ B_GC_ cells than ED-primed RMs at week 4 (median 14.8% vs. 1.3%, *P* = 0.0004; **Fig. 1G**). Although the frequency of IgM^+^ cells declined in both groups by week 8, it remained significantly higher in bolus-primed RMs (*P* < 0.0001). Clonal diversity of the Env⁺ GC response was also assessed. Slow-delivery priming recruited more than 10 times as many Env^+^ B_GC_ clones as bolus priming by week 4 (median 266 vs. 12.5 clones, *P* < 0.0001; **Fig. 1H**). Although clone numbers in both groups declined over time, they remained significantly higher in ED-primed RMs at week 8 (*P* = 0.0065). Chao1 richness estimates and the Gini– Simpson diversity index similarly demonstrated greater repertoire diversity after slow-delivery priming (**Fig. 1I-J**).

The broader repertoire induced by ED priming suggested recruitment of B cell clones spanning a wider range of antigen affinities, potentially including low-affinity precursors that fall below the detection even of tetravalent probe binding. To quantify the recruitment of very low-affinity precursors, Env⁻ LN B cells isolated at week 4 were sequenced and clonally linked by VDJ sequencing to Env^+^ cells. Thus, although these cells were Env⁻ by flow cytometry at week 4, their clonal relationship to Env⁺ cells identified them as members of Env-specific lineages that initially fell below the detection threshold of the Env probe. A median of 29% of Env⁻ B_GC_ clones from ED-primed RMs were clonally related to Env⁺ cells, significantly more than bolus-primed animals (*P* = 0.0012), with only three bolus-primed RMs exhibiting any Env⁻ to Env⁺ clonal overlap (**Fig. 1K**). These data support the hypothesis that a slow-delivery prime recruits a broader repertoire of low-affinity B cell clones than bolus priming, and those low-affinity clones subsequently undergo substantial affinity maturation.

If slow delivery lowers the effective affinity threshold for B cell recruitment, it could also enable recruitment of clones recognizing subdominant Env epitopes that would otherwise be excluded because of low initial affinity. To determine whether this broader recruitment extended to subdominant Env specificities, responses to the highly immunodominant non-glycosylated base of the Env trimer were assessed^40–42^. A base-knockout (BaseKO) probe, which is not easily recognized by base-specific Env⁺ B_GC_ cells, was used to identify BaseKO⁺ Env^+^ B_GC_ cells, representing cells recognizing non-base Env epitopes (**Fig. S1G**). Despite the broader recruitment observed following slow delivery, the size of the non-base response did not differ between bolus- and ED-primed RMs (**Fig. S1H**).

Together, these data establish that slow-delivery priming generates larger, more proliferative, and more clonally diverse GC responses than bolus immunization, accompanied by accelerated class switching and enhanced recruitment and affinity maturation of low-affinity B cell clones. However, this broader early recruitment was not accompanied by a detectable increase in responses to subdominant non-base Env epitopes.

### Env⁺ GCs persist for nearly a year after a slow-delivery prime

Having established the early differences between bolus and slow-delivery priming, we next asked how long a single prime could sustain GC activity. Previous work demonstrated that Env^+^ GCs remained detectable in the draining LN for six months following a single ED prime^16^. Here, Group 4 remained unboosted until week 52, allowing us to follow the prime-only GC for up to one year and determine whether GC activity could persist beyond 6 months.

Env^+^ B_GC_ cells were quantified longitudinally in LN FNA samples by flow cytometry across all four groups (**Fig. 2A; Fig. S2A**). In the ED-primed groups, GC responses expanded to a peak around week 10 and subsequently declined but remained detectable through week 48 in the absence of boosting. In contrast, bolus priming generated substantially smaller primary GC responses, with robust GC expansion occurring only after boosting. Following boosting, both bolus groups generated robust GC responses that remained readily detectable through week 48, consistent with persistence for approximately six months after the week 24 boost. Comparison of peak GC magnitude showed that the pooled ED-primed groups had significantly larger responses than Group 1 (P = 0.0098; **Fig. S2B**), whereas the response in Group 2 was not significantly different from either Group 1 or the ED-primed groups. Notably, GC responses in Groups 1 and 2 showed similar persistence following the week 24 boost, despite Group 1 having received an additional boost at week 10 (**Fig. 2A)**.

**Figure 2.**
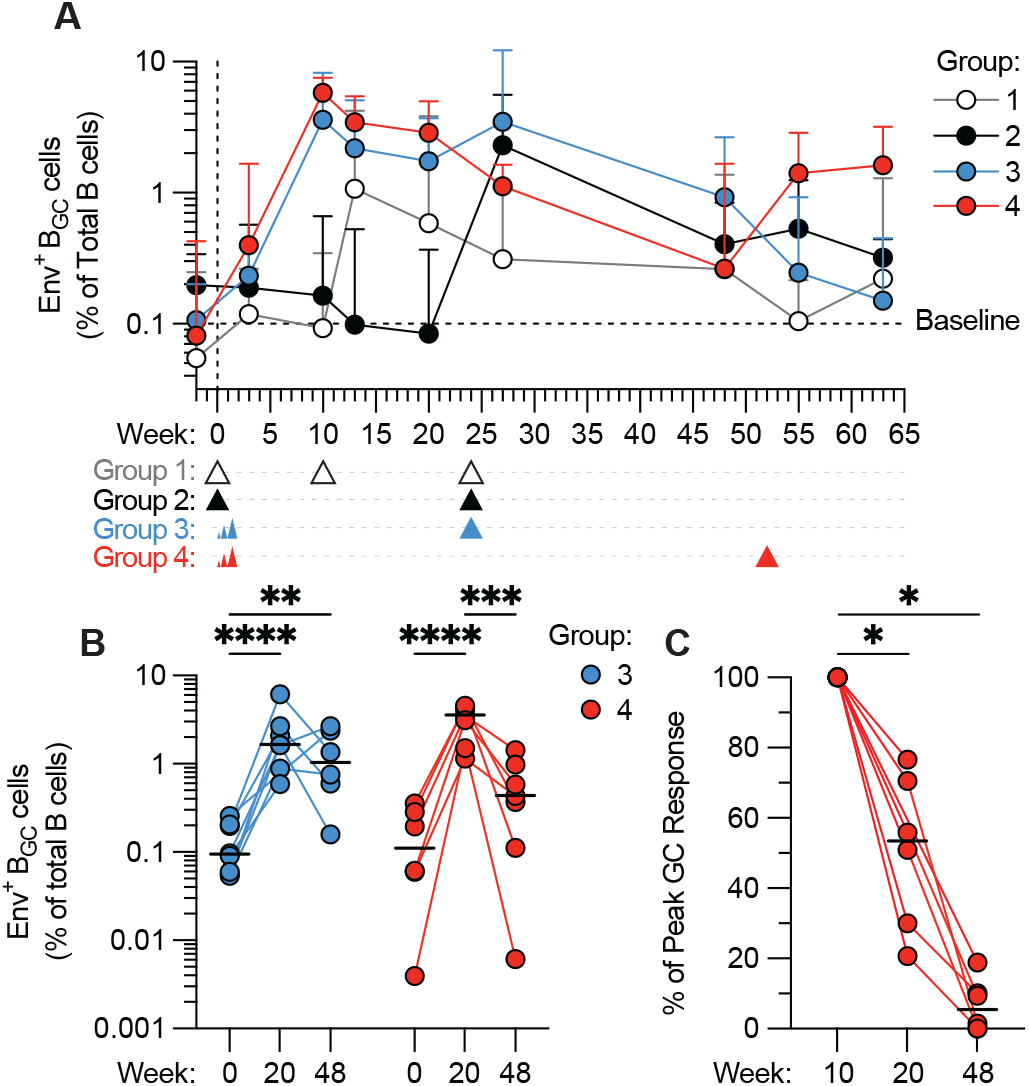
A single ED prime sustains Env⁺ GC responses for up to 48 weeks. **(A)** Frequency of Env⁺ B_GC_ cells as a percentage of total B cells in LN FNAs over time. The horizontal dotted line indicates the median baseline frequency (week −2), and the vertical dotted line indicates immunization at week 0. Arrows below the x-axis indicate booster immunizations; single arrows denote bolus immunization, and a series of increasing arrows denotes escalating-dose (ED) immunization. Error bars indicate mean ± standard error of the mean (SEM). **(B)** Frequency of Env⁺ B_GC_ cells as a percentage of total B cells in ED-primed animals (Groups 3 and 4) at baseline (week −2), week 20 and week 48. **(C)** Persistence of the Env⁺ B_GC_ cell response in Group 4 animals at weeks 20 and 48, expressed relative to each animal’s peak week-10 response after baseline correction (baseline = 0%; week 10 = 100%). Statistical comparisons were performed using a mixed-effects analysis with multiple-comparisons correction for **(B)** and two-tailed Wilcoxon matched-pairs signed-rank tests for **(C)**. P values are indicated as P < 0.05 (*), P < 0.01 (**), P < 0.001 (***), and P < 0.0001 (****); non-significant comparisons are not shown. Horizontal bars indicate median.

To quantify Env^+^ GC persistence more directly, GC size at baseline (week −2) was compared to weeks 20 and 48 in the ED-primed groups (**Fig. 2B**). At week 20, both groups reflected the response to ED priming alone, and Env^+^ B_GC_ responses remained significantly above baseline (Group 3, *P* < 0.0001; Group 4, *P* < 0.0001), demonstrating sustained GC activity for at least five months following priming. Thereafter, Group 3 received a boost at week 24, whereas Group 4 remained unboosted. At week 48, Env⁺ GC responses in Group 3 remained significantly above baseline (median 1.02% at week 48 vs. 0.09% at baseline; *P* = 0.013), consistent with the week-24 boost sustaining an elevated response for at least a further 24 weeks. In contrast, Env^+^ B_GC_ frequencies in the unboosted Group 4 were no longer significantly different from baseline (median 0.43% at week 48 vs. 0.11% at baseline; *P* = 0.21; **Fig. 2B**).

The extent of primary Env^+^ GC contraction was then assessed in the unboosted Group 4 by normalizing each animal’s response to its own peak at week 10 (**Fig. 2C**). GC size declined significantly between weeks 10 and 20 (*P* = 0.031) and decreased further by week 48 (*P* = 0.031), at which point all but one animal exhibited less than 10% of their peak Env^+^ GC activity. Thus, primary GC responses had substantially contracted by one year, suggesting that 48 weeks may approach the upper limit of primary GC persistence in this model. Interestingly, a similar analysis of Group 3 showed that Env^+^ GC responses appeared to have fully resolved by the end of the study, despite boosting at six months (**Fig. S2C**).

Together, these data demonstrate that a single ED prime can sustain Env^+^ GCs for close to a year without further antigen exposure. In contrast, bolus priming generated substantially smaller primary GC responses, but a booster immunization resulted in sustained GC activity for an additional six months. However, additional boosting appeared to only modestly increased GC persistence.

### Long-lived GCs remain clonally dynamic and progressively shift toward subdominant Env epitopes

To examine how GC repertoires evolved in these long-lived GC responses, individual Env⁺ B_GC_ clones were tracked longitudinally by paired VDJ sequencing. GC repertoires remained dynamic throughout the response, with substantial changes in the relative representation of sampled clones during both the priming phase and following booster immunization (**Fig. 3A**). Clones were classified as detected during priming (prime-only), boosting (boost-only), or both periods (prime+boost). Only a small fraction of clones fell into the prime+boost category, but these persistent clones accounted for a disproportionately large fraction of Env^+^ B_GC_ cells (**Fig. 3B-C**) and were on average significantly larger than either prime-only or boost-only clones (**Fig. S3A**).

**Figure 3.**
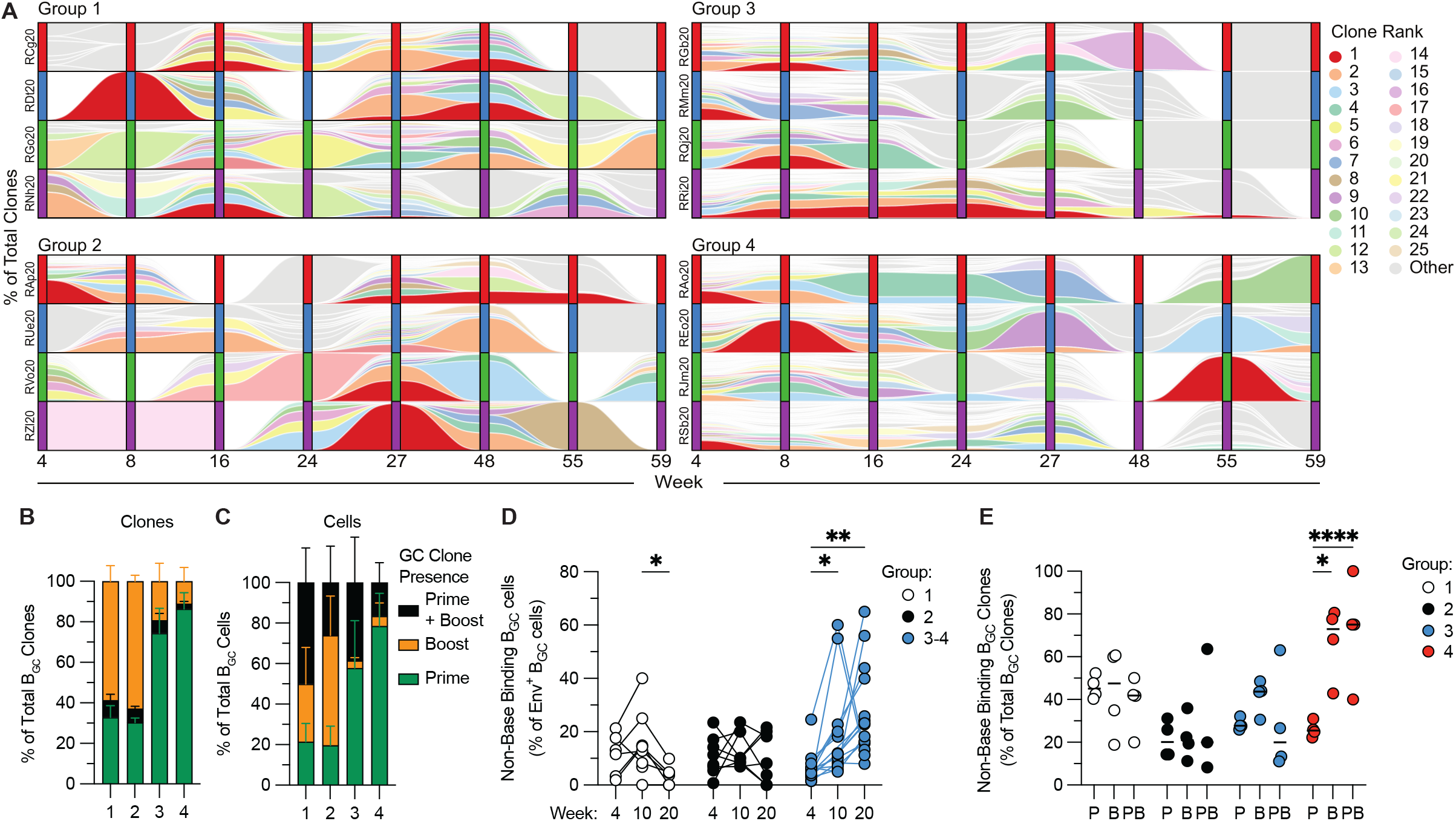
Long-lived germinal centers undergo continual clonal turnover and progressive shifts in epitope specificity. **(A)** Alluvial plots showing longitudinal clonal dynamics of Env⁺ B_GC_ cells. At each time point, clone abundance is shown as a proportion of the total Env⁺ B_GC_ repertoire for each animal per group. The 25 largest clones in each animal are colored and tracked over time; all remaining clones are shown in gray. White space indicates time points at which a clone was not detected. **(B)** Distribution of Env⁺ B_GC_ clones according to whether they were detected during priming only (Prime), following boosting only (Boost), or during both periods (Prime + Boost). **(C)** Distribution of Env⁺ B_GC_ cells belonging to Prime, Boost, or Prime + Boost clones as defined in **(B)**. **(D)** Frequency of non-base-binding B_GC_ cells among Env⁺ B_GC_ cells during the priming period (weeks 4, 10, and 24), shown separately for Groups 1 and 2 and the combined ED-primed groups (Groups 3 and 4). **(E)** Frequency of non-base-binding B_GC_ clones among Prime (P), Boost (B), and Prime + Boost (PB) clones. Statistical comparisons were performed using two-way ANOVA with multiple-comparisons correction for **(D)** and **(E)**. P values are indicated as P < 0.05 (*), P < 0.01 (**), P < 0.001 (***), and P < 0.0001 (****); non-significant comparisons are not shown. Horizontal bars indicate median.

This dynamic clonal composition contrasted with the more stable clonal composition of the antigen-specific T follicular helper (Tfh) response previously characterized in the same animals^17^, consistent with continued reshaping of the B cell response during affinity maturation. We therefore asked whether the epitope specificities within the Env⁺ GC changed over time. Longitudinal analysis of BaseKO probe binding revealed that the proportion of non-base-binding Env^+^ B_GC_ cells increased significantly between weeks 3 and 10 in ED-primed animals (P = 0.029) and remained elevated through week 20 (median 4.1% at week 3 vs. 23% at week 20, P = 0.0022; **Fig. 3D; Fig. S3B**), demonstrating progressive enrichment for B_GC_ cells targeting subdominant Env epitopes. No comparable increase was observed following bolus priming.

To determine whether this shift reflected changes in clonal composition, clones were classified as non-base-binding according to the proportion of non-base-binding cells they contained. In Group 4, the frequency of non-base-binding clones was significantly higher among both boost-only (P = 0.0005) and prime + boost (P = 0.0001) clones than among prime-only clones following the delayed week-52 boost (**Fig. 3E**). Interestingly, this shift in clonal composition was unique to Group 4, despite Groups 3 and 4 having comparable post-boost frequencies of non-base-binding Env^+^ B_GC_ cells by flow cytometry, implying that delayed boosting broadened the clonal composition of the non-base response without increasing its overall magnitude (**Fig. S3C**).

### Somatic hypermutation rapidly decelerates after the first months of the GC response

We next examined the accumulation of SHM in Env^+^ B_GC_ cells over time. Although SHM is the molecular engine of affinity maturation, how quickly mutations accumulate over the lifetime of a GC has rarely been quantified because it requires dense longitudinal sampling of the same response over many months. The depth of longitudinal LN FNA sampling in this study allowed for direct examination of SHM rates. Heavy-chain (HC) nucleotide (nt) mutations per Env^+^ B_GC_ cell were quantified over time (**Fig. 4A-B**). Mutation loads increased progressively throughout the response. Notably, SHM accumulation was nearly identical across all four groups (**Fig. 4A**), including the bolus groups, which had significantly smaller Env^+^ GC responses following priming (**Fig. 1C, 2A**). These substantial mutation loads could only have accumulated through prolonged residence within active GCs, demonstrating that the smaller responses elicited by bolus priming nevertheless supported sustained GC evolution for at least 6 months post prime. B_Mem_ cells and B_PCs_ isolated from the LNs showed the same progressive accumulation of SHM, with slightly lower median mutation loads at each time point, consistent with these populations comprising recent GC emigrants that had ceased accumulating mutations (**Fig. S4A-B**).

**Figure 4.**
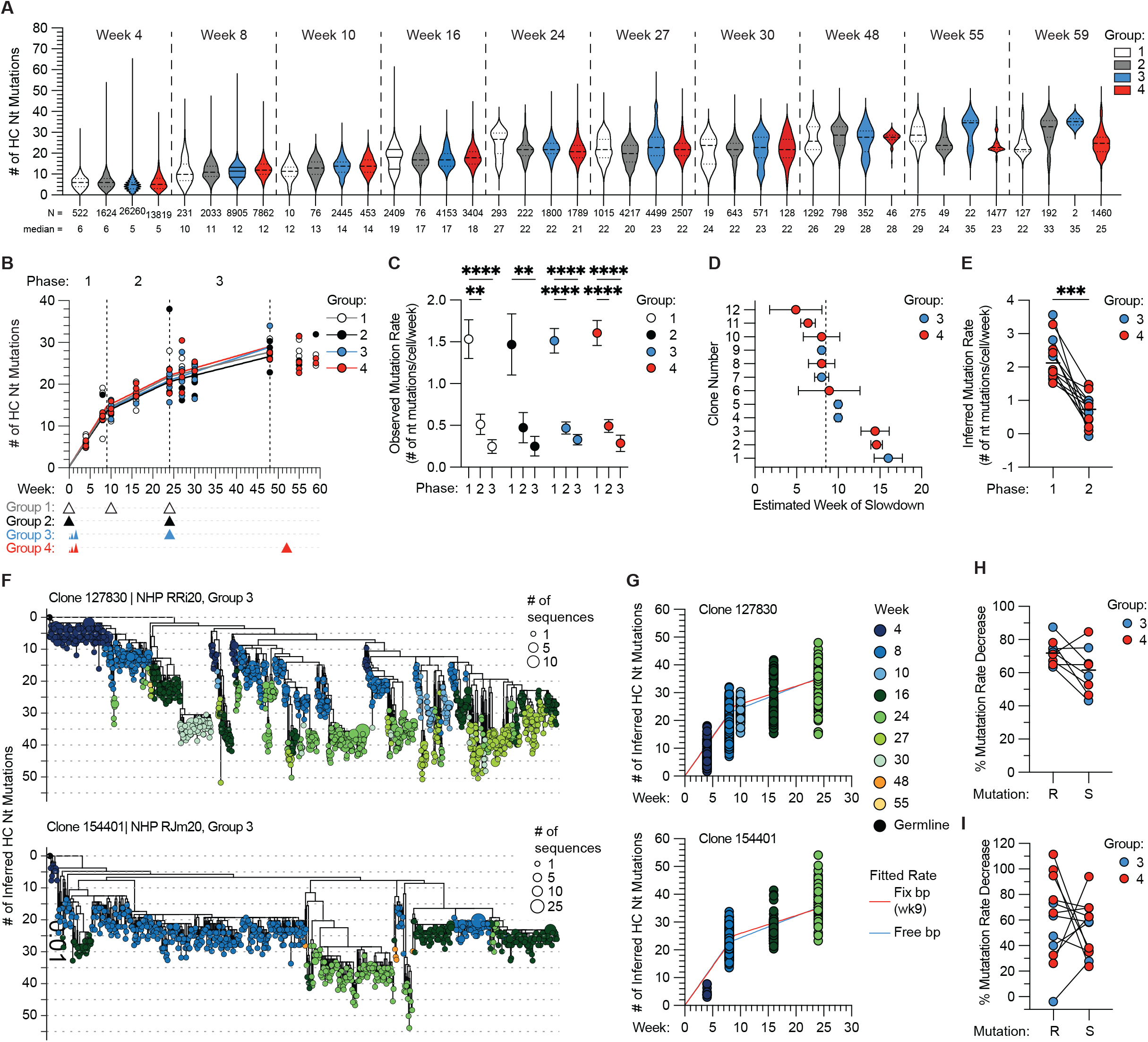
The rate of somatic hypermutation in Env⁺ B_GC_ cells declines sharply early in the GC response. **(A)** Heavy-chain (HC) nucleotide (Nt) mutations per Env⁺ B_GC_ cell in all groups over time. Horizontal lines indicate the median and quartiles. The number of cells (N) and median mutation count are shown below each violin. **(B)** Mean HC Nt mutations per animal over time, colored by group. Lines show the fitted three-phase trajectory from a piecewise linear mixed-effects model fit to per-animal weekly means, with each animal–week contributing equally. Breakpoints were set at week 9, determined by segmented regression of the pooled data, and week 24, corresponding to the boost given at that time point for Groups 1, 2, and 3. Vertical dotted lines mark the phase boundaries: phase 1 (weeks 0–9), phase 2 (weeks 9–24), and phase 3 (weeks 24–48). Arrows below the x-axis indicate booster immunizations. **(C)** Observed mutation rate (HC Nt mutations per B_GC_ cell per week) for each group in each phase, derived from the model in (B). Error bars indicate 95% confidence intervals. **(D)** Estimated breakpoint (week of slowdown) for the twelve persistent clones colored by group. Dotted line indicates the median week of slowdown (8.5). Error bars indicate 95% confidence intervals. **(E)** Inferred mutation rate per clone in phases 1 and 2, colored by group; lines connect phases within a clone. Inferred mutations from germline were estimated from maximum-likelihood clonal trees, accounting for repeat and back mutation at the same site. Per-clone rates were obtained by piecewise regression of inferred mutations from germline against sampling week, with the breakpoint set at week 9. **(F)** Clonal lineage trees for two clones spanning multiple time points. Dotted lines indicate 5-mutation intervals; the y-axis shows inferred mutations from germline. Tips are colored by the time point at which each cell was collected, and tip size is proportional to sequence count. **(G)** Inferred mutations from germline per cell over time for the clones in (F). Lines denote fitted mutation rates, red when the breakpoint was set at week 9 and blue when estimated independently for that clone. **(H)** Percentage decrease in observed mutation rate from phase 1 to phase 2 for replacement (R) and silent (S) mutations in all ED-primed animals, colored by group. **(I)** Percentage decrease in observed mutation rate from phase 1 to phase 2 for replacement (R) and silent (S) mutations in persistent clones, colored by group. Statistical comparisons were performed within the mixed-effects model for **(C),** with phase 1 versus phase 2 compared directly from the corresponding hinge coefficient and phase 1 versus phase 3 as a linear combination of both hinge terms; and using the Wilcoxon matched-pairs signed-rank test for **(E)**, **(H)**, and **(I)**. P values are indicated as P < 0.05 (*), P < 0.01 (**), P < 0.001 (***), and P < 0.0001 (****); non-significant comparisons are not shown. Horizontal bars indicate median.

Inspection of the mutation trajectories suggested that SHM accumulation slowed as the response progressed (**Fig. 4A-B**). To determine when this transition occurred, mutation load was modeled as a function of time using sequences pooled from Groups 2–4, whose priming period through week 24 was uninterrupted by boosting. A segmented regression model allowing a single change in slope fit the data significantly better than a constant-rate model (Davies test, P = 1.6 × 10⁻⁶) and identified a breakpoint, marking the slowdown in mutation accumulation, at week 9.2 ± 1.3. Mutation accumulation rates were therefore estimated across three phases using a linear mixed-effects model: Phase 1, before the slowdown (weeks 0–9); Phase 2, from the slowdown to week 24 (weeks 9–24); and Phase 3, following the week-24 boost (weeks 24–48), with Group 4 remaining unboosted throughout this interval (**Fig. 4B**). Later time points were excluded due to limited sampling depth.

Mutations accumulated rapidly during the first phase in all groups (1.47–1.61 nt mutations/cell/week) before slowing approximately three-fold during the second phase (0.47–0.51 nt mutations/cell/week; *P* < 0.0001 to 0.004, and *P* = 0.060 in Group 2) (**Fig. 4C**). Rates declined further during the third phase (0.25–0.33 nt mutations/cell/week), representing a 4.6- to 6.2-fold reduction from the first phase (P = 0.0013 to P < 0.0001), although the third phase did not differ significantly from the second in any group (Fig. 4C). Strikingly, phase-specific mutation accumulation rates were nearly identical across all four groups (group × phase interaction, P = 0.93). Bolus- and ED-primed animals accumulated mutations at indistinguishable rates (first-phase difference −0.03 nt mutations/cell/week, 95% CI −0.31 to 0.24). Similarly, second-phase rates did not differ between Group 1, boosted at week 10, and Group 2, which remained unboosted (difference −0.03, 95% CI −0.44 to 0.39), or between boosted Group 3 and unboosted Group 4 during the third phase (difference −0.05, 95% CI −0.27 to 0.18). Together, these comparisons demonstrate that neither priming strategy nor antigen re-exposure measurably altered the rate of mutation accumulation within the Env^+^ GC response (**Fig. 4C**).

Because the preceding analyses pooled sequences across many independent B cell clones, we next asked whether the same slowdown could be observed within individual clonal lineages. Twelve clones meeting a predefined sampling threshold of at least five cells across two or more time points in each of Phase 1 and Phase 2 were selected for phylogenetic reconstruction. Root-to-tip path length was used to estimate mutations accumulated along each lineage. Because these distances are inferred from reconstructed phylogenies, the resulting rates are on a different scale from the per-cell/animal rates (**Fig. 4A-C)**. Applying the same segmented regression approach to each clone independently identified breakpoints, representing the estimated week of slowdown, ranging from week 4.9 to week 16, with a median of week 8.5 (**Fig. 4D**), closely matching the repertoire-wide breakpoint of week 9.2. Thus, the population-level slowdown likely reflects an average across lineages that slow at different times rather than a synchronized transition across the repertoire. Inferred mutation rates declined significantly from the first to the second phase within these clones (median 2.12 vs. 0.73 inferred mutations/cell/week, *P* = 0.0005; **Fig. 4E**), recapitulating the population-level slowdown. Two deeply sampled clones illustrate these dynamics, with estimated breakpoints at week 10 for clone 127830 and week 8 for clone 154401 compared with the population-level breakpoint at week 9 (**Fig. 4F-G**).

To determine whether these findings generalized across studies, the same analyses were applied to an independent cohort of ED-primed macaques from a previously published study^16^. A similar breakpoint was identified at week 7.9 ± 1.2 (**Fig. S4C**), with mutation accumulation slowing 4.3-fold from 1.71 to 0.40 mutations/cell/week (*P* < 0.0001; **Fig. S4D**). Clone-level analysis again recapitulated this pattern, with inferred mutation rates declining from 2.22 during phase 1 (≤ week 8) to 0.79 mutations/cell/week during phase 2 (> week 8) (*P* = 0.0012; **Fig. S4E**) and a median clonal breakpoint at week 7.1 (**Fig. S4F**).

The striking consistency of this slowdown across groups and studies suggested a common underlying mechanism. One possibility is increasing affinity-dependent selection. As affinity matures, an increasing fraction of replacement mutations may become detrimental to antigen binding and be removed by selection, whereas phenotypically neutral silent mutations should continue to accumulate relatively unaffected. To test this, silent and replacement mutation accumulation was compared in the ED groups (Groups 3 and 4) across the first two phases, where sequencing depth was greatest. At the repertoire level, both mutation types slowed to a comparable degree between phases 1 and 2 (P = 0.078), although replacement mutations showed a slightly greater reduction in five of eight animals (**Fig. 4H**). Comparable slowing of replacement and silent mutation accumulation was also observed within persistent clones (P = 0.62; **Fig. 4I**) and in the independent cohort (**Fig. S4G**). The comparable deceleration of silent and replacement mutations at both the repertoire and clonal levels argues that the slowdown is not primarily explained by increasing affinity-dependent selection.

Another potential explanation is a progressive decline in the intrinsic mutability of B cell receptor (BCR) sequences. Activation-induced cytidine deaminase (AID) targets nucleotides unevenly^43^. Preferred hotspots may therefore be progressively consumed as SHM proceeds, reducing the number of mutations each cycle can generate^44^. However, sequences remained more mutable than their germline precursors throughout the response, with median mutability declining only from 1.07 to 1.03 between phases (P = 0.0078; ∼4%), far too little to account for the observed three-fold reduction in rate (**Fig. S4H**).

Finally, the slowdown could potentially reflect changes in GC cellular dynamics that reduce opportunities for continued SHM as the response matures. To investigate this, GC cellular composition was examined over the course of the response (**Fig. S4I**). The proportion of proliferating B_GC_ cells declined over time, while the LZ:DZ ratio increased through week 24 (**Fig. S4J-K**). This shift was driven in part by a specific increase in MYC^-^ LZ cells (**Fig. S4L**). Because MYC induction in LZ B cells is associated with receipt of Tfh help and subsequent re-entry into the proliferative DZ program^45–47^, the accumulation of MYC^-^ LZ cells may indicate that a progressively smaller fraction of LZ cells undergoes productive selection as GCs age.

Together, these data demonstrate a pronounced transition from rapid to slower SHM accumulation during prolonged GC evolution. This mutational deceleration was reproducible across independent cohorts, was independent of priming strategy and boosting and was recapitulated within individual clonal lineages, indicating that mutational deceleration represents an intrinsic feature of long-lived GC evolution.

### Slow-delivery priming generates a durable circulating memory compartment enriched for B cells targeting subdominant Env epitopes

Circulating B_Mem_ cells are a major output of the GC. To determine how prolonged GC evolution shaped the circulating memory compartment, Env^+^ B_Mem_ frequencies in peripheral blood were tracked longitudinally across all groups (**Fig. S5A**). At week 10, slow-delivery priming induced ∼10-fold higher Env^+^ B_Mem_ frequencies than bolus priming (median 0.29% vs. 0.03%, *P* < 0.0001; **Fig. 5A**). Booster immunization induced a marked but transient expansion of Env^+^ B_Mem_ cells in all groups, reaching approximately 1–3% of IgD⁻ B cells two weeks post boost before contracting to a resting frequency of approximately 0.1–0.2% (**Fig. 5B**). Following ED priming alone, Env^+^ B_Mem_ frequencies in Group 4 gradually declined after their week 10 peak (**Fig. 5B**; **Fig. S5C**) but remained detectable after one year.

**Figure 5.**
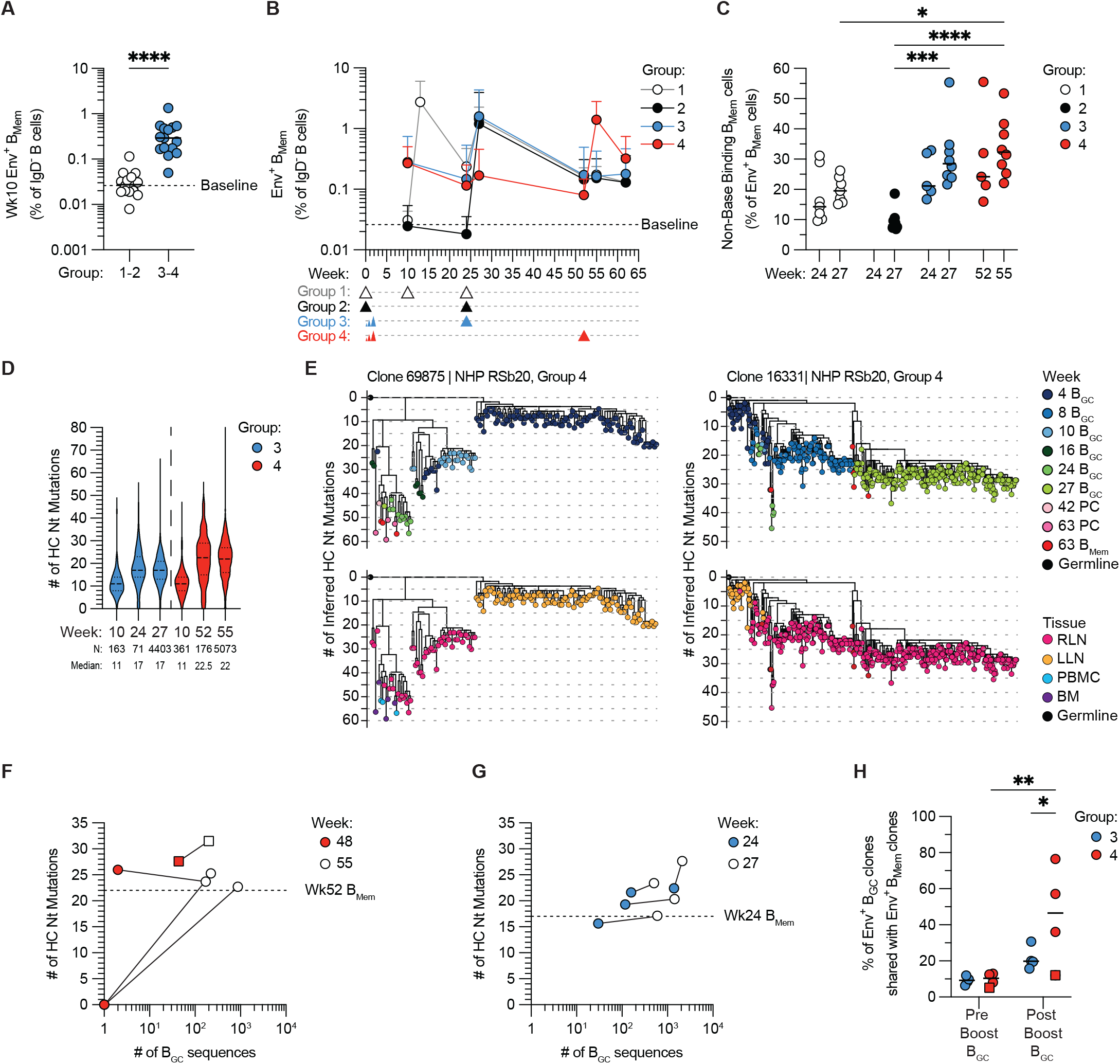
Prolonged GC responses sustain a persistent and highly mutated circulating Env⁺ B_Mem_ compartment that can contribute to secondary GC responses. **(A)** Frequency of circulating Env⁺ B_Mem_ cells as a percentage of IgD⁻ B cells at week 10 following bolus (Groups 1 and 2) or ED (Groups 3 and 4) priming. The horizontal dotted line indicates the median baseline frequency (week −2), calculated from Groups 1 and 3. **(B)** Frequency of circulating Env⁺ B_Mem_ cells as a percentage of IgD⁻ B cells over time. Error bars indicate mean ± SEM. Dotted line indicates the baseline frequency, calculated from Groups 1 and 3. Arrows below the x-axis indicate booster immunizations. **(C)** Frequency of non-base-binding B_Mem_ cells among circulating Env⁺ B_Mem_ cells before and after boosting. Group 2 at week 24 is not shown because too few Env⁺ B_Mem_ cells were recovered to reliably estimate the non-base-binding fraction. Pre-boost measurements are unavailable for three animals each in Groups 3 and 4 because of limited sample availability. **(D)** Heavy-chain (HC) nucleotide (Nt) mutations per Env⁺ B_Mem_ cell in ED-primed animals over time. Horizontal lines indicate the median and quartiles. Cell numbers (N) and median mutation counts are shown below each violin. **(E)** Clonal lineage trees for two representative clones detected in both draining LNs. Tips are colored by sampling time point (top) or tissue of origin (bottom). The y-axis indicates the number of inferred HC Nt mutations from germline; dotted horizontal lines indicate five-mutation intervals. **(F)** Relationship between the number of Env⁺ B_GC_ sequences recovered and median HC Nt mutations in Group 4 animals before (week 48) and after (week 55) boosting. Paired samples from individual animals are connected by lines. The dotted line indicates the median HC Nt mutation count of circulating Env⁺ B_Mem_ cells in Group 4 before boosting (week 52). The animal with a clearly detectable pre-boost GC response is denoted by square symbols. Samples with no recovered B_GC_ sequences are plotted at 1 sequence for visualization on the logarithmic x-axis and assigned a mutation count of zero. **(G)** Relationship between the number of Env⁺ B_GC_ sequences recovered and median HC Nt mutations in Group 3 animals before (week 24) and after (week 27) boosting. Paired samples from individual animals are connected by lines. The dotted line indicates the median HC Nt mutation count of circulating Env⁺ B_Mem_ cells in group 3 before boosting (week 24). **(H)** Percentage of Env⁺ B_GC_ clones sampled before or after boosting that were clonally related to circulating Env⁺ B_Mem_ clones. The animal with a clearly detectable pre-boost GC response is denoted by a square symbol. Statistical comparisons were performed using a two-tailed Mann–Whitney test for **(A)** and two-way ANOVA with multiple-comparisons correction for **(C)** and **(H)**. P values are indicated as P < 0.05 (*), P < 0.01 (**), P < 0.001 (***), and P < 0.0001 (****); non-significant comparisons are not shown. Horizontal bars indicate median.

Although Env^+^ B_Mem_ frequencies were comparable after boosting, the specificity of the memory compartment differed between immunization regimens. Reflecting the progressive shift toward subdominant specificities in GCs (**Fig. 3D; Fig. S3B**), non-base-binding Env^+^ B_Mem_ frequencies were generally higher following slow delivery than bolus priming after boosting (**Fig. 5C**). Extending the boost interval from six months to one year did not significantly increase the size of the non-base B_Mem_ response between the two ED-primed groups.

### Evidence for B_Mem_ recirculation through prolonged GCs

To understand how the memory compartment evolved over the course of the prolonged GC response, paired VDJ sequencing was performed on sorted Env^+^ B_Mem_ cells collected at week 10, immediately before boosting, and after boosting in the ED-primed groups. As expected, circulating Env^+^ B_Mem_ cells accumulated SHM over time (**Fig. 5D**). However, this increase was accompanied by a marked loss of the lowly mutated B_Mem_ cells abundant at week 10. Approximately 40–45% of Env^+^ B_Mem_ cells carried ≤10 HC nt mutations at week 10, compared with only ∼8.5–11% immediately before boosting in Groups 3 and 4 (**Fig. S5D**). This loss of lowly mutated B_Mem_ cells occurred despite a gradual decline, rather than expansion, of the total circulating Env^+^ B_Mem_ population over the same period (**Fig. 5B; Fig. S5C**), arguing against their simple dilution by increasingly mutated B_Mem_ cells.

One possible explanation for this loss is that Env^+^ B_Mem_ cells generated earlier in the response re-enter ongoing GCs and undergo further SHM. We previously observed that a subset of GC clonal lineages could be detected in both draining LNs following bilateral slow-delivery priming^16^, consistent with recirculation of B_Mem_ cell clones between anatomically distinct GC responses. To determine whether this phenomenon was also observed here, clonal lineage trees were reconstructed from paired VDJ sequences obtained from both draining lymph nodes, with longitudinal sampling of the LLN and a single week 4 sample from the RLN. Multiple clonal lineages were again detected in GCs from both draining LNs (**Fig. 5E**), with approximately 1.5% of all observed B_GC_ clones shared between LNs (**Table S1**). Although lower than the ∼5% overlap observed previously^16^, this was not unexpected given that bilateral LN sampling in the current study was limited to a single time point, whereas both draining LNs were sampled longitudinally in the previous study.

### B_Mem_ cells contribute to secondary GC responses

The potential for B_Mem_ cells to participate in ongoing GC responses raises the broader question of how the memory compartment contributes to GCs following antigen re-exposure. B_Mem_ cells can participate in recall responses by differentiating into antibody-secreting cells (ASCs) or reseeding secondary GCs, although the extent of their contribution to secondary GCs remains debated^48–50^. The different boost timings in Groups 3 and 4 provided an opportunity to examine B_Mem_ recruitment into secondary GCs across substantially different levels of pre-existing GC activity, with median Env^+^ B_GC_ frequencies at the last sampled pre-boost time point of 1.62% in Group 3 compared with 0.43% in Group 4 (**Fig. 2B**).

Given that circulating B_Mem_ cells were slightly less mutated than contemporaneous B_GC_ cells (**Fig. 4A**; **Fig. 5D**), their recruitment in the GC following boosting might be expected to reduce the mutation load of the post-boost GC population. In Group 4, where Env^+^ GCs had largely contracted before the week-52 boost, median HC nt mutation counts decreased from 28 at week 48 to 23 at week 55 (**Fig. 4A**), closely matching the median of 22.5 mutations in circulating Env^+^ B_Mem_ cells immediately before boosting (**Fig. 5D**). This decrease was not accompanied by an increase in very lowly mutated Env⁺ B_GC_ cells, suggesting limited recruitment of naive B cells into the Env-binding GC response following boosting. This relationship was also evident at the individual-animal level: animals with absent or very small pre-boost GCs showed a post-boost decrease in GC mutation levels that approached those of pre-boost B_Mem_ cells, whereas a single animal with a clearly detectable pre-existing GC showed an increase in mutation levels (**Fig. 5F**). In contrast, following the earlier boost in Group 3, when Env^+^ GCs were still large, mutation levels increased in all animals and generally remained higher than those of pre-boost B_Mem_ cells (**Fig. 5G**).

To test whether this shift was also reflected in clonal composition, Env^+^ B_GC_ repertoires sampled before and after boosting were compared based on the percentage of clones also detected among that animal’s circulating Env^+^ B_Mem_ compartment. Because too few pre-boost B_Mem_ cells were available for a direct comparison with the post-boost GC, B_Mem_ sequences were pooled across all time points, with the resulting repertoire dominated by post-boost samples. Following the delayed week-52 boost, a substantially greater proportion of Group 4 Env^+^ B_GC_ clones sampled after boosting were also detected among circulating B_Mem_ clones compared with those sampled before boosting (median 46.4% vs. 10.3%, P = 0.0095; **Fig. 5H**). No significant increase was observed following the earlier week-24 boost in Group 3 (P = 0.24). Notably, the Group 4 animal with the lowest post-boost overlap was also the animal with the largest residual pre-boost GC response and the same animal in which B_GC_ mutation levels increased following boosting. Because the pooled B_Mem_ repertoire was dominated by post-boost cells, this increased overlap could reflect B_Mem_ cells generated from the GC following the boost rather than B_Mem_ recruitment into it. To assess this possibility, the converse relationship was examined to ask whether clones present in the GC before boosting were subsequently detected in the post-boost B_Mem_ repertoire. This overlap was low (∼10%) and did not differ between Groups 3 and 4 (**Fig. S5E**), suggesting that the much greater B_GC_–B_Mem_ overlap observed in Group 4 was not driven by increased GC-to-B_Mem_ output.

Together, the reduction in GC mutation levels toward those of circulating B_Mem_ cells and the nearly five-fold increase in clonal overlap between the GC and B_Mem_ compartments support a model in which secondary GCs formed by boosting after the primary GC has largely contracted are increasingly seeded by circulating B_Mem_ cells. In contrast, when boosting occurs while the primary GC is still ongoing, existing B_GC_ clones may make a proportionally greater contribution to the response.

### Delayed boosting broadens antibody specificity without enhancing neutralization

The circulating antibody response was next examined to assess how prolonged GC evolution was reflected at the serological level. Slow-delivery priming elicited robust Env-specific serum IgG titers after the primary immunization, while a bolus prime generated little detectable Env-specific serum IgG until after boosting (**Fig. 6A**). Following boosting, differences in resting Env-binding antibody titers between groups were substantially reduced despite their markedly different post-prime responses (**Fig. S6A**), consistent with prior findings^21,36^.

**Figure 6.**
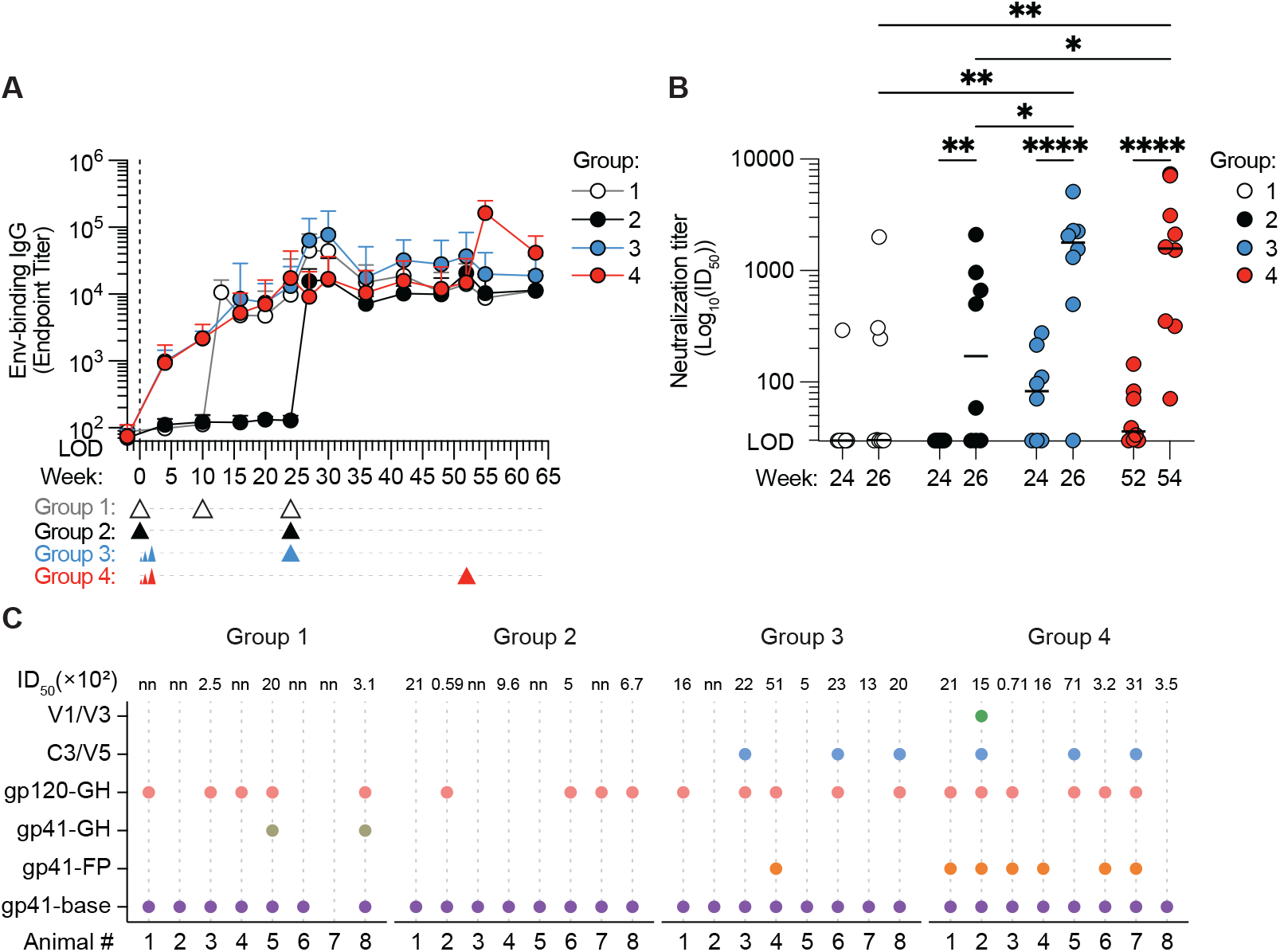
Delayed boosting broadens epitope targeting without improving neutralizing antibody titers. **(A)** Env-specific serum IgG endpoint titers over time, colored by group. Error bars indicate mean ± SEM. The y-axis begins at the limit of detection (LOD), and the vertical dotted line indicates immunization at week 0. Arrows below the x-axis indicate booster immunizations. **(B)** Autologous neutralizing antibody titers (50% inhibitory dilution; ID_50_) before and two weeks after boosting, colored by group. **(C)** Serum antibody epitope specificities determined by electron microscopy polyclonal epitope mapping (EMPEM) two weeks after boosting, shown for all eight animals in each group. Colored circles indicate detection of antibody responses targeting the indicated HIV Env epitopes, grouped by their location within gp41 or gp120. Autologous ID_50_ values (×10²) measured two weeks post-boost are shown above each animal; nn denotes non-neutralizing. Statistical significance in **(B)** was determined using two-way ANOVA with multiple-comparisons correction. P values are indicated as P < 0.05 (*), P < 0.01 (**), P < 0.001 (***), and P < 0.0001 (****); non-significant comparisons are not shown. Horizontal bars indicate median.

Although all groups ultimately generated comparable Env-binding serum antibody titers, much of the Env-specific antibody response is directed against the non-neutralizing base epitope^41,42^. Binding titers therefore may not reflect the induction of neutralizing antibodies. To directly assess neutralizing activity, autologous neutralizing antibody responses were measured (**Fig. 6B; Fig. S6B**). Prior to boosting, small but detectable autologous neutralizing antibody titers were observed in 8 of 16 ED-primed animals, recapitulating our previous finding that ED priming alone can elicit neutralizing antibodies, whereas only 1 of 16 bolus-primed animals had detectable neutralizing activity (Fisher’s exact test, P = 0.0155). Following boosting, detectable neutralizing responses were observed in 15 of 16 ED-primed animals compared with 8 of 16 bolus-primed animals (Fisher’s exact test, P = 0.0155), and neutralizing antibody titers were significantly higher in both ED groups than in either bolus group (Group 1 vs. Group 3, P = 0.0011; Group 1 vs. Group 4, P = 0.0016; Group 2 vs. Group 3, P = 0.0343; Group 2 vs. Group 4, P = 0.0463; **Fig. 6B**), confirming the superior immunogenicity of ED priming compared with bolus priming. However, delaying the boost from six months to one year did not further increase neutralizing antibody titers between groups 3 and 4.

Given the progressive enrichment of B_GC_ cell responses for subdominant Env epitopes over time, we next asked whether delayed boosting nevertheless altered the specificity of the circulating antibody response. Electron Microscopy Polyclonal Epitope Mapping (EMPEM)^42^ was performed on serum collected two weeks after boosting. Nearly all RMs generated antibodies targeting the immunodominant Env base, whereas responses against several subdominant epitopes were observed predominantly in ED-primed animals (**Fig. 6C; Fig. S6C-D**). Group 4 also exhibited the highest overall EMPEM magnitude (average number of antibodies per immune complex) (**Fig. S6D**), consistent with its higher Env-binding IgG endpoint titers at the post-boost time point (**Fig. 6A; Fig. S6A**). Most notably, fusion peptide-proximal antibodies were detected in 6 of 8 Group 4 animals but in only a single animal across all other groups, indicating that delayed boosting broadened the circulating antibody response to include additional subdominant Env epitopes. Post-boost serum neutralization titers were also highest in ED-primed animals, consistent with the detection of antibodies targeting V1/V3, C3/V5, and gp41-FP, epitopes previously associated with autologous neutralization in NHPs immunized with BG505-based Env immunogens^51,52^ (**Fig. S6B-D**).

### Ongoing BM B_PC_ generation during prolonged GC responses is modulated by booster timing

As the source of circulating antibody, B_PCs_ represent the major effector output of the GC. We therefore sought to determine how the Env^+^ BM B_PC_ compartment developed over the course of these long-lived GC responses. To track the establishment and evolution of this compartment over time, BM aspirates were collected at weeks 24, 30, and 42, and femoral BM scoops obtained at necropsy (week 63). Env^+^ B_PCs_ were quantified by ELISpot as IgG^+^ Env^+^ ASCs relative to total IgG⁺ ASCs. At week 24, before boosting, ED-primed RMs had a readily detectable population of Env⁺ ASCs in the BM (**Fig. 7A-B**). Env⁺ ASCs were ∼14-fold more frequent following ED priming than a single bolus prime (Group 2; median 1.1% vs. 0.08%, *P* = 0.0001) but were not significantly different from Group 1, which had received its first boost at week 10 (*P* = 0.65; **Fig. 7A**). All three groups boosted at week 24 (Groups 1–3) showed a significant increase in Env⁺ ASCs by six weeks after boosting, and these elevated responses remained relatively stable through to necropsy at week 63 (**Fig. 7B**). Group 4 remained unchanged between weeks 24 and 42 but underwent a similarly robust expansion following the booster immunization at week 52 (**Fig. 7B; Fig. S7B**).

**Figure 7.**
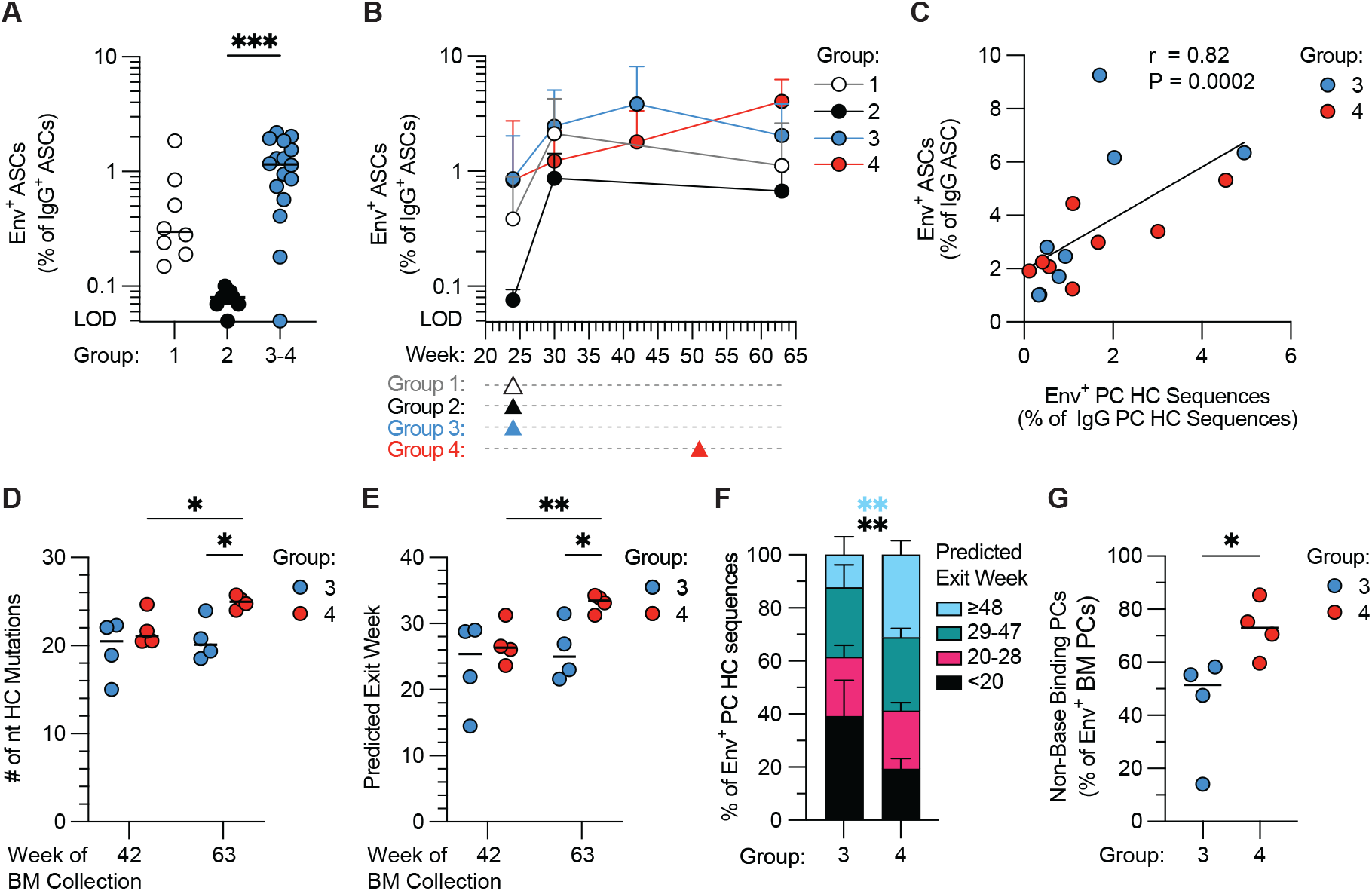
Delayed boosting enriches the bone marrow (BM) B_PC_ compartment for late GC emigrants and subdominant Env specificities. **(A)** Env⁺ antibody-secreting cells (ASCs) as a percentage of total IgG⁺ ASCs in BM at week 24, shown for Group 1, Group 2, and the combined ED-primed groups (Groups 3 and 4). The y-axis begins at the limit of detection (LOD). **(B)** Env⁺ ASCs as a percentage of total IgG⁺ ASCs in BM over time, colored by group. Error bars indicate mean ± SEM. The y-axis begins at the LOD. Arrows below the x-axis indicate booster immunizations. **(C)** Correlation between Env⁺ ASCs as a percentage of total IgG⁺ ASCs, determined by ELISpot, and Env⁺ B_PC_ HC sequences identified by clonal overlap with Env⁺ B_GC_ or B_Mem_ cells, as a percentage of total IgG B_PC_ HC sequences. *r* indicates the Spearman correlation coefficient. **(D)** Mean number of HC Nt mutations per Env⁺ BM B_PC_ sequence for each animal in samples collected at weeks 42 and 63, colored by group. **(E)** Mean predicted GC exit week of Env⁺ BM B_PC_ sequences for each animal in samples collected at weeks 42 and 63, colored by group. **(F)** Distribution of Env⁺ BM B_PC_ HC sequences predicted to have exited the GC before week 20, between weeks 20– 28, between weeks 29–47, or at or after week 48, shown by group in week-63 BM samples. Bars indicate mean ± SEM. *P* values are colored according to the corresponding predicted exit interval. **(G)** Frequency of non-base-binding B_PCs_ among Env⁺ BM B_PCs_ at week 63, colored by group. Horizontal bars indicate medians. Statistical comparisons were performed using a Kruskal–Wallis test with multiple-comparisons correction for **(A),** Spearman correlation for **(C)**, two-way ANOVA with multiple-comparisons correction for **(D)** and **(E)**, and two-tailed Mann–Whitney tests for **(F)** and **(G)**. P values are indicated as P < 0.05 (*), P < 0.01 (**), P < 0.001 (***), and P < 0.0001 (****); non-significant comparisons are not shown. Horizontal bars indicate median.

The apparent stability of the Env^+^ BM B_PC_ compartment observed over time could reflect the establishment of a long-lived plasma cell (LLPC) population. However, because GCs remain active for up to a year after priming, this stability could instead be maintained by the continual production of new B_PCs_ exiting the GC. Boosting may further alter this process by driving additional B_PC_ generation. Distinguishing between these possibilities requires knowing when individual B_PCs_ were generated, which cannot currently be directly determined outside of genetic fate-mapping approaches in engineered mouse models. We therefore sought to estimate when BM B_PCs_ exited the GC using SHM as a molecular clock. Having established that B_GC_ cells accumulate SHM at a predictable rate, albeit one that slows over time (**Fig. 4**), we reasoned that a B_PC’s_ mutational load could be used to estimate when it exited the GC and ceased accumulating mutations. These analyses focused on the slow-delivery-primed groups, allowing B_PC_ generation to be examined across prolonged GC responses and following boosting at different stages of GC evolution.

To estimate GC exit times, Env^+^ BM B_PCs_ first needed to be identified and sequenced, but because B_PCs_ downregulate surface BCR, they cannot be readily sorted using fluorescent Env probes. Instead, we exploited the extensive reference repertoire of Env^+^ B_GC_ and B_Mem_ cells generated throughout the study. B_PC_ HC VDJ sequences belonging to clones represented in this repertoire could be identified as Env-specific by matching VDJs of Env^+^ B_GC_ and B_Mem_ cells. Bulk HC VDJ sequencing was performed on sorted CD20^low^CD38^+^ BM B_PC_s at weeks 42 and 63, sorting both CD19^+^ and CD19⁻ B_PCs_ at weeks 42 and 63 (**Fig. S7B**). Env⁺ B_PC_ sequences were then identified by clonal matching to the Env^+^ B_GC_ and B_Mem_ repertoires. The frequency of clonally matched Env^+^ B_PC_ sequences strongly correlated with the frequency of Env⁺ ASCs measured by ELISpot (Spearman r = 0.82, P = 0.0002; **Fig. 7C; Fig. S7C**), validating clonal matching as an approach for identifying bona fide Env⁺ B_PCs_. Having identified Env⁺ B_PCs_ by clonal relationship, we examined their SHM levels as an initial measure of when these cells were generated (**Fig. 7D**). Because CD19 expression has been associated with potentially less mature BM B_PC_ populations^53^, we first asked whether CD19^+^ and CD19⁻ Env^+^ B_PCs_ differed in their mutation load as a potential indicator of relative B_PC_ age. At week 63, HC mutation loads were similar between CD19⁺ and CD19⁻ Env⁺ BM B_PC_s in Groups 3 and 4 (**Fig. S7D**). However, given the limited number of CD19^+^ sequences and animals available for this comparison, subsequent analyses considered the BM B_PC_ compartment as a whole, regardless of CD19 expression. Across the combined BM B_PC_ compartment, the mutation loads of B_PCs_ collected at week 42 were comparable between Groups 3 and 4 (**Fig. 7D**). Following the delayed week-52 boost, mutation levels of B_PCs_ collected at week 63 in Group 4 increased significantly (*P* = 0.016) and exceeded those in Group 3 (median 25 vs. 20 mutations; P = 0.024). In contrast, mutation levels in Group 3 remained unchanged over time.

To place these differences in SHM on a temporal scale, each B_PC_ mutation count was mapped onto the group-specific mutation-versus-time trajectory determined in **Fig. 4B–C** to estimate when that level of SHM would have been reached within the GC. Because the mutation-rate model extended only to week 48, estimated GC exit times beyond week 48 were capped at this value. At the week 42 and week 63 BM collection time points, median estimated GC exit in Group 3 remained around week 25, close to the timing of the week-24 boost (**Fig. 7E**). In contrast, the median estimated GC exit time in Group 4 shifted from approximately week 25 to week 32 (*P* = 0.0081), resulting in significantly later estimated GC exit time than Group 3 at the week 63 collection time point (*P* = 0.037; **Fig. 7E**). B_PC_ sequences were next grouped into blocks of estimated GC exit time (<week 20, weeks 20–28, weeks 29–47, or ≥week 48; **Fig. 7F**) to determine which periods of B_PC_ generation accounted for this shift. B_PCs_ were represented across all four intervals in both groups, indicating continued generation of BM B_PCs_ throughout the prolonged GC response. Surprisingly, despite Group 3 receiving its boost at week 24, the proportion of B_PCs_ with an estimated GC exit between weeks 20–28 was nearly identical between groups. Instead, Group 3 contained a significantly greater proportion of B_PCs_ with an estimated GC exit before week 20 (mean 39.2% vs. 19.3%, *P* = 0.0019), while significantly fewer had an estimated GC exit at or after week 48 (mean 12.3% vs. 31%, *P* = 0.0034; **Fig. 7F**).

The greater contribution of late GC emigrants to the Group 4 B_PC_ compartment raised the possibility that this compartment would also be enriched for the subdominant non-base Env specificities that progressively emerged during GC maturation (**Fig. 3**), potentially providing a cellular basis for the broader serum epitope recognition observed following delayed boosting (**Fig. 6C**). Using the same BaseKO classification applied to B_GC_ clones, Env^+^ B_PC_ sequences were classified as base-binding or non-base-binding. Group 4 contained a significantly greater proportion of non-base-binding B_PCs_ than Group 3 (*P* = 0.0286; **Fig. 7G**), demonstrating that the greater representation of late GC emigrants following delayed boosting was accompanied by increased representation of more subdominant epitope specificities in the BM B_PC_ compartment.

A similar analysis of the bolus-primed groups at the week-63 BM collection indicated that B_PCs_ generated throughout the prolonged GC response contributed to the BM compartment in both groups, with no significant differences in the inferred timing of B_PC_ generation between Groups 1 and 2 (**Fig. S7E-I**). Although Group 1 received an additional boost at week 10, both groups received their final boost at week 24, potentially limiting the ability to resolve an effect of the earlier boost at this later time point.

Collectively, these findings demonstrate that a slow-delivery prime is sufficient to establish an Env^+^ B_PC_ population in the BM. B_PCs_ contributing to this compartment were generated throughout the prolonged GC response, rather than being restricted to a particular stage of GC maturation. Booster timing nevertheless left a lasting imprint on its composition, with earlier boosting associated with greater representation of early-generated B_PCs_ and delayed boosting increasing the contribution of the most mature GC output. This shift enriched the BM B_PC_ repertoire for subdominant non-base Env specificities.

## DISCUSSION

The recognition that vaccine-induced GCs can persist for many months has substantially expanded the timescale over which B cell affinity maturation is thought to occur. However, previous longitudinal studies have generally ended while antigen-specific GCs remained active, leaving both their ultimate lifespan and capacity for continued evolution unresolved. In this study, extended longitudinal analysis revealed that vaccine-induced GCs can remain active and continue to evolve up to a year, with ongoing SHM and major impacts on the dynamics and quality of both B_Mem_ and BM B_PC_ compartments.

Env^+^ B_GC_ cells remained detectable 48 weeks after a slow-delivery prime, extending the known duration of a primary vaccine-induced GC response beyond the six months previously observed. GC responses were substantially diminished by this time, indicating that one year may approach the upper limit of primary GC persistence in this model. Notably, long-lived GCs were not unique to slow delivery priming: despite much smaller responses following bolus priming, continued SHM and prolonged post-boost GCs demonstrated that extended GC maturation can also follow conventional immunization. Interestingly, additional boosting augmented ongoing responses but did not appear to reset their lifespan, suggesting that GC longevity is not simply determined by renewed antigen exposure. Together with similar observations in humans following both infection and immunization^11,13–15,18^, these findings demonstrate that long-lived GCs are likely a general feature of robust humoral responses.

One presumed benefit of long-lived GCs is continued SHM and affinity maturation, yet here it was observed that the rate of SHM accumulation slowed markedly over time, falling from ∼1.5 to ∼0.3–0.5 observed HC nt mutations per B_GC_ cell per week. Consequently, approximately as much SHM accumulated during the first nine weeks as over the subsequent thirty-nine weeks. Understanding what drives this slowdown requires considering what SHM rate represents mechanistically. Classical estimates place SHM at ∼10⁻³ mutations per base pair per cell division or ∼0.35 mutations per HC V(D)J sequence^48,49^. However, recent studies indicate that SHM is largely deferred to a post-mitotic window at the end of the DZ proliferative burst, allowing multiple divisions to occur without additional mutation^50,51^. Thus, ∼0.35 HC mutations may better approximate mutations per productive GC cycle than per cell division, raising the possibility that SHM accumulation provides a molecular readout of productive GC recycling. Under this framework, the early SHM rate corresponds to ∼4 productive GC cycles per week, or approximately one cycle every two days. Although this may be somewhat slower than the rapid GC cycling rates inferred from mouse studies^1,4,54^, the difference could reflect slower GC dynamics in NHPs. By contrast, the later rate corresponds to just ∼1 productive cycle per week, which would imply substantial slowing of GC recycling as the response matures.

Interpreting slower SHM accumulation as reduced GC recycling, however, assumes that the number of mutations acquired and retained per cycle remains relatively constant. The slowdown could instead result from fewer mutations generated per cycle or increased removal of newly mutated cells through selection. Neither increased selection nor declining sequence-intrinsic mutability, however, appeared to account for the slowdown. Although other mechanisms affecting mutational yield cannot be excluded, these data strengthen the possibility that productive GC recycling itself becomes less frequent as the response matures.

This raises the question of which stage of the GC cycle might account for the slowing in SHM. One possibility is that changes in LZ dynamics increasingly limit the pace of GC recycling. The increasing proportion of MYC⁻ LZ cells over time suggest that longer GC cycles could reflect increased time spent in the LZ. B_GC_ cells must acquire antigen and compete for Tfh help in the LZ before positively selected cells can re-enter the DZ and undergo further proliferation and SHM^1,4,46,55^. As GCs mature and the availability of accessible antigen declines, LZ cells may require more time to acquire sufficient antigen and Tfh help for DZ re-entry. This could increase LZ dwell time and thereby extend the interval between successive rounds of SHM. However, the relationship between antigen and Tfh availability and the pace of GC recycling is unlikely to be straightforward. Despite substantial differences in antigen availability and Tfh responses between slow-delivery and bolus priming^17,21^, SHM slowed at remarkably similar times and to similar rates, and boosting did not restore rapid SHM. Thus, although altered LZ dynamics provide a plausible explanation for slower productive GC recycling, how antigen availability, Tfh help, B cell competition, and other features of the maturing GC interact to potentially determine the pace of recycling remains unresolved.

The failure of boosting to restore rapid SHM may reflect the fact that the earlier boosts appeared to primarily refuel ongoing GCs rather than initiate new responses. When boosting instead occurred as the primary GC response approached its end, however, there was evidence of substantial recruitment of recalled B_Mem_ cells, consistent with the formation of de novo secondary GCs. SHM kinetics could not be reliably estimated following this late boost, leaving open whether substantial B_Mem_ recall can reset SHM accumulation toward rates observed early in the response. Determining whether newly initiated secondary GCs undergo a renewed period of rapid SHM will be important for understanding how affinity maturation proceeds across repeated antigen exposures. This question has broad implications for vaccination, including sequential immunization strategies designed to iteratively mature specific B cell lineages and recurrent vaccination against evolving pathogens such as influenza virus, where the capacity of recalled B cells to rapidly acquire new mutations may influence adaptation to antigenically distinct strains. More broadly, these findings suggest that booster timing may influence whether antigen re-exposure predominantly sustains an existing GC response or reseeds secondary GCs from the memory compartment. The optimal interval may therefore depend on whether continued selection of B_GC_ clones already participating in the response or broader recruitment from the evolved B_Mem_ compartment is desired.

Importantly, the B_Mem_ compartment itself continued to evolve during prolonged GC activity. Although circulating Env⁺ B_Mem_ cells accumulated SHM over time, this was accompanied by a disproportionate loss of the lowly mutated B_Mem_ cells abundant early in the response, consistent with similar evolution of the B_Mem_ repertoire observed following SARS-CoV-2 infection and vaccination^56–58^. Because the overall Env⁺ B_Mem_ compartment contracted rather than expanded over this period, this shift is unlikely to reflect simple dilution by newly generated GC emigrants. Instead, these observations suggest continual turnover of the circulating memory compartment. One possibility is that early lowly mutated B_Mem_ cells are progressively lost over time. Alternatively, these cells could re-enter ongoing GCs and undergo further diversification, with both processes potentially contributing to the observed changes. Consistent with the possibility of GC re-entry, GC clones were detected across anatomically distinct draining lymph nodes before boosting, both here and in our previous study^16^, suggesting that B cell clones can recirculate between GC responses, likely through a B_Mem_ intermediate. Although only ∼1.5–5% of B_GC_ clones were detected in both draining LNs, this likely underestimates the extent of such recycling, as re-entry into GCs within the same LN would not be distinguishable from continued participation in the original GC response in this model. Thus, these data suggest that the memory compartment may remain dynamically coupled to ongoing GC evolution rather than representing a static archive of earlier stages of the response.

Finally, a principal function of the GC is the generation of high-affinity, long-lived B_PCs_ that provide durable protection through sustained antibody production. How the long-lived B_PC_ compartment is established over the course of a GC response lasting many months, however, remains unclear. Several models could explain how this compartment is populated during such prolonged responses, including continuous B_PC_ generation throughout GC maturation, preferential B_PC_ differentiation during later stages of the response, or increased B_PC_ output following antigen re-exposure^29–33^. Distinguishing these models is challenging because there are no definitive markers of B_PC_ age. However, the predictable accumulation of SHM observed here provided a means to approximate when BM B_PCs_ diverged from the evolving GC repertoire. This analysis suggests that the BM B_PC_ compartment is populated throughout prolonged GC maturation, with little evidence that B_PC_ differentiation or accumulation becomes intrinsically favored as GCs age. Instead, booster timing appeared to alter the relative contribution of early- and late-generated cells to the BM B_PC_ compartment. This has important implications for sequential vaccination, where the goal is not simply to generate increasingly mature B cell lineages, but ultimately to capture those matured states within the durable antibody-producing compartment. For HIV vaccination, delaying a final “polishing” immunization^25^ until bnAb lineages have sufficiently matured may therefore help preferentially recruit these cells into the persistent BM B_PC_ compartment, translating their maturation into durable antibody production.

One caveat to this interpretation is that mutation-based dating for B_PCs_ cannot distinguish direct B_GC_ differentiation from differentiation of recalled B_Mem_ cells. Because B_Mem_ cells in our study were generally less mutated than contemporaneous B_GC_ cells, B_Mem_-derived B_PCs_ could be assigned to somewhat earlier stages of the response. Nevertheless, because B_Mem_ SHM broadly tracked that of B_GC_ cells over time, mutation frequency should still provide a reasonable estimate of the general stage of the response from which these B_PCs_ were derived.

Together, these findings demonstrate that prolonged GCs remain dynamic long after their initial expansion, with continued evolution of epitope specificity and downstream B cell responses despite progressively slower SHM. Rather than simply extending the early phase of GC maturation, long-lived GCs therefore represent a distinct phase of the response, whose changing properties should be considered when designing vaccination strategies that rely on prolonged and iterative affinity maturation.

### Limitations of the Study

Although this study provides an extensive longitudinal characterization of vaccine-induced GC responses, two limitations should be acknowledged. First, while the study included 32 RMs, longitudinal single-cell RNA and VDJ sequencing was performed on only four animals per group, limiting statistical power for comparisons of clonal dynamics and repertoire composition. Second, LN FNA sampling captures only a fraction of the total GC response, meaning that individual clones may be missed at particular time points. Consequently, estimates of clonal persistence and overlap between GC and B_Mem_ compartments likely underestimate the extent of clonal continuity and recycling.

## Data availability

Single-cell sequencing data generated in this study can be accessed in the Gene Expression Omnibus database under accession number (pending). Representative EMPEM reconstructions have been deposited to the Electron Microscopy Data Bank (EMDB) under accession codes EMD-79405, EMD-79406, EMD-79412 and EMD-79414. Any additional information required to reanalyze the data reported in this paper is available from the lead contact upon request.

## Acknowledgments

We thank the staff of the Emory National Biomedical Research for their extensive support with animal care, immunizations, and sample collection throughout this study. We are especially grateful to the Flow Cytometry and Next Generation Sequencing core facilities at the La Jolla Institute for Immunology for their invaluable technical expertise and support. We also thank members of the Crotty laboratory for critical reading of the manuscript and helpful discussions. This work was supported in part by the National Institute of Allergy and Infectious Diseases of the NIH under award numbers CHAVD UM1AI144462 and R01 AI125068, and by the Emory National Biomedical Research through ORIP/OD grants P51OD011132 and U42 PDP11023.

## Contributions

Conceptualization, H.J.S. and S.C.; Methodology, H.J.S., E.M.Z., D.G.C., D.J.I., and S.C.; Formal Analysis, H.J.S., T.K.A., A.P., A.A.U., I.B., M.B., and G.O.; Investigation, H.J.S., E.M.Z., T.K.A., A.P., A.A.U., I.B., M.B., W.-H.L., K.N.R.J., A.S.T., K.H., B.S.H., and D.L.; Resources, A.W., M.B.M., J.T., N.P., D.Lu., D.G., C.A.C., W.R.S., D.J.I., and S.C.; Writing – Original Draft, H.J.S. and S.C.; Writing – Review & Editing, H.J.S., E.M.Z., D.G.C., L.H., D.R.B., G.S., A.B.W., G.O., S.E.B., S.P.K., W.R.S., D.J.I., and S.C.; Project Administration, H.J.S., D.G.C., and S.C.; Supervision, L.H., D.R.B., D.G.C., G.S., C.A.C., A.B.W., G.O., S.E.B., S.P.K., W.R.S., D.J.I., and S.C.; Funding Acquisition, D.R.B., G.S., W.R.S., D.J.I., and S.C.

## METHODS

### Protein and SMNP Expression

BG505 MD39 SOSIP Env trimers (MD39) were co-expressed with furin in HEK293F or Expi293 cells as previously described^59^. Trimers used for immunizations, ELISPOT, and EMPEM were made without an affinity tag, purified using 2G12 affinity chromatography followed by size exclusion chromatography (SEC), and quality checked for low endotoxin levels as previously described^60^. BG505 MD39 SOSIP and BG505 MD39-base knockout trimers (BaseKO) used as flow cytometry probes and in ELISAs were generated with C-terminal His and Avi-tags, purified using immobilized metal ion affinity chromatography followed by SEC, and biotinylated using a BirA biotinylation kit according to the manufacturer’s instructions (Avidity, catalog no. BirA500) as previously described^60^. The BG505 MD39-base knockout trimer had the following mutations relative to the BG505 MD39 SOSIP: A73C, R500A, P561C, C605T, S613T, Q658T, L660N, A662T and L663C. SMNP adjuvant was prepared and characterized as previously described^34^

### Animals, immunizations and sample acquisition

The animal cohort, immunization regimens, and sample collection schedule used in this study have been described previously in a study of the T follicular helper cell response in the same animals^17^. Briefly, Indian rhesus macaques (RMs; *Macaca mulatta*) were housed at the Emory National Biomedical Research Center and maintained in accordance with NIH guidelines. All procedures were approved by the Emory University Institutional Animal Care and Use Committee (IACUC) under protocol 202100121. Animal care facilities are accredited by the U.S. Department of Agriculture (USDA) and the Association for Assessment and Accreditation of Laboratory Animal Care (AAALAC) International.

Animals were distributed among four study groups (n = 8 per group), balanced for age, weight, and sex. RMs were 2.4–2.7 years of age at the time of the initial immunization. Immunizations were administered subcutaneously and divided equally between the left and right mid-thighs. Each immunization contained a total of 100 μg MD39 Env trimer and 750 μg saponin/MPLA nanoparticle (SMNP) adjuvant, corresponding to 50 μg MD39 and 375 μg SMNP per injection site. The four immunization regimens were: Group 1, bolus prime followed by boosts at weeks 10 and 24; Group 2, bolus prime followed by a boost at week 24; Group 3, escalating-dose (ED) prime followed by a boost at week 24; and Group 4, ED prime followed by a boost at week 52.

For Groups 3 and 4, the priming immunization was administered using the previously described 12-day ED regimen^16,21^. Seven injections were given every other day from days 0–12, with antigen and adjuvant amounts increasing exponentially over the dosing period. MD39 was administered at 0.1, 0.215, 0.58, 1.575, 4.28, 11.65, and 31.6 μg per side, and SMNP at 0.75, 1.1625, 4.35, 11.81, 32.1, 87.375, and 237 μg per side. All booster immunizations were delivered as single bolus injections using the same immunogen, adjuvant, and total dose as the bolus priming immunization. Thus, the cumulative amount of MD39 and SMNP administered per immunization was equivalent between bolus and ED regimens.

Fine-needle aspirates (FNAs) were collected longitudinally from the left and right inguinal lymph nodes (iLNs). Draining LNs were identified by palpation, and a 22-gauge needle attached to a 3-mL syringe was passed through each LN up to five times. Aspirated cells were transferred into RPMI containing 10% fetal bovine serum (FBS) and 1× penicillin/streptomycin. Samples with visible red blood cell contamination were treated with ammonium-chloride-potassium (ACK) lysis buffer. Cells were subsequently counted and resuspended in freezing medium containing 10% DMSO in FBS. Because FNA cell yields varied between collections, samples were cryopreserved at the available cell concentration rather than normalized to a fixed concentration. Samples were frozen at −80°C and subsequently transferred to liquid nitrogen for long-term storage.

Peripheral blood was collected into sodium citrate CPT tubes (BD Biosciences) for isolation of peripheral blood mononuclear cells (PBMCs) and plasma. PBMCs were isolated according to the manufacturer’s instructions, counted, and resuspended in freezing medium containing 10% DMSO in FBS at approximately 1×10^7^ cells/mL. Cells were cryopreserved at −80°C and subsequently transferred to liquid nitrogen for long-term storage. Plasma was aliquoted and stored frozen until analysis. Serum was collected using serum-separator/clot tubes and stored frozen until analysis.

Bone marrow (BM) was collected longitudinally by aspiration and at necropsy by femoral scoop. For BM aspirates, 1–3 mL of marrow was collected into heparinized syringes, diluted in PBS, and layered over lymphocyte separation medium (LSM) at a 2:1 sample-to-LSM ratio. Samples were centrifuged at 1452×g for 30 min without brake. The mononuclear cell layer was collected, washed in PBS, and centrifuged at 600×g for 10 min. Residual erythrocytes were removed by ACK lysis for 5 min at room temperature, followed by washing in PBS. Cells were resuspended in complete medium and counted by trypan blue exclusion using a Countess automated cell counter. Cells were then resuspended in freezing medium containing 10% DMSO in FBS at approximately 1×10^7^ cells/mL, cryopreserved at −80°C, and subsequently transferred to liquid nitrogen for long-term storage.

At necropsy, femoral BM was mechanically dissociated through a 70–100-µm cell strainer using PBS and the base of a 5-mL syringe. The resulting cell suspension was filtered to remove residual tissue and debris, underlaid with Ficoll, and centrifuged at 1452×g for 30 min without brake. The mononuclear cell layer was collected, washed in PBS, and treated with ACK lysis buffer for 5 min at room temperature to remove residual erythrocytes. Cells were then washed, resuspended in complete medium, and counted. Femoral BM cells were cryopreserved in 10% DMSO in FBS at approximately 1×10^7^ cells/mL and stored in liquid nitrogen.

Animals received anesthesia (ketamine, 5–10 mg/kg, or Telazol, 3–6 mg/kg) and appropriate analgesia for immunizations, LN FNAs, bone marrow aspirates and blood collections according to veterinary recommendations and IACUC-approved procedures.

### Flow cytometry and cell sorting

Frozen FNA and PBMC samples were thawed in R10 medium (RPMI containing L-glutamine and HEPES, supplemented with 10% FBS, 1% penicillin/streptomycin, and 1% GlutaMAX). Cells were centrifuged at 500×g for 5 min at 37°C, resuspended in R10 medium, and counted. Following counting, cells were washed, resuspended in fluorescence-activated cell sorting (FACS) buffer (PBS containing 2% FBS), and transferred to 96-well plates for staining.

Fluorescent MD39 WT and MD39 knockout (KO) probes were generated by conjugating biotinylated antigen to fluorophore-conjugated streptavidin at a 2:1 mass ratio. One-third of the total streptavidin mixture was added to the biotinylated probe every 15 min over 45 min. Probe conjugates were diluted in PBS such that 0.5 μg of antigen was delivered per 5×10^6^ cells. Cells were first incubated with MD39 KO probe at 2.5 μL per 5 × 10^6^ cells for 15 min at 4°C, followed by two differently fluorophore-labeled MD39 WT probes, each added at 2.5 μL per 5 × 10^6^ cells, for an additional 15 min at 4°C. Probe volumes were scaled proportionally for samples containing larger cell numbers. Cells were then stained with the appropriate phenotyping antibody panel (**Table S2**) for a further 30 min at 4°C. For cell sorting, sample-specific TotalSeq-C hashtag antibodies (BioLegend) were added together with the phenotyping panel at 2 μg per 5×10^6^ cells.

Following staining, cells were washed twice with FACS buffer and resuspended for acquisition or sorting. FNA samples were sorted on either a FACSymphony S6 or FACSAria Fusion (BD Biosciences), whereas PBMC samples were sorted on a FACSymphony S6. Flow cytometric analysis was performed on a 5-laser Cytek Aurora (Cytek Biosciences). For FNA samples, a minimum of 250 total B cells and 75 GC B cells was required for inclusion in flow cytometric analyses. Samples below either threshold were considered unsuccessful FNAs and excluded from analysis.

Bone marrow B_PCs_ were stained and sorted by flow cytometry. Approximately 10–15×10^6^ bone marrow mononuclear cells (BMMCs) were used for BM B_PC_ staining and sorting. Cells were first stained with eBioscience™ Fixable Viability Dye eFluor™ 780 (Cat#65-0865-14, Thermo Fisher Scientific) at a 1:5000 dilution in PBS for 30 min at 4 °C to exclude dead cells. Following washing, Fc receptors were blocked using anti-Rh Fc binding inhibitor (Cat#14-9165-42, Invitrogen), and cells were surface stained for 30 min at 4 °C in FACS buffer (1× PBS supplemented with 2% FBS and 2 mM EDTA) with the appropriate phenotyping antibody panel (**Table S2**). Following surface staining, cells were washed, resuspended in FACS buffer, and sorted in 350 µL of RLT buffer (QIAGEN) using a BD FACSAria II (BD Biosciences). All flow cytometry data were analyzed in FlowJo v10 software (BD Biosciences).

### Single-cell RNA library preparation and sequencing

Sorted B cells were loaded onto Chromium Next GEM Single Cell 5′ v2 chips for FNA samples or v3 chips for PBMC samples and processed using the Chromium Controller according to the manufacturer’s instructions (10x Genomics). Gene expression (GEX), V(D)J, and Feature Barcode libraries were generated using the Dual Index 10x Genomics Single Cell 5′ V(D)J with Feature Barcode workflow. Custom primers targeting rhesus macaque BCR constant regions were included at final concentrations of 1 μM for forward primers and 0.5 μM for reverse primers. Libraries were pooled and sequenced on an Illumina NovaSeq 6000.

### BM B_PC_ adaptive immune receptor repertoire sequencing (AIRR-seq) library preparation and sequencing

AIRR-seq libraries were prepared using a previously described method developed by the Daniel Douek laboratory at the National Institute of Allergy and Infectious Diseases (NIAID) Vaccine Research Center^61,62^. RNA was extracted from sorted BM B_PCs_ using the RNeasy Micro kit with DNase digestion (QIAGEN) on a QIAcube automated platform. cDNA was generated using SMARTer template switching (Clontech). RNA was combined with a 5′ CDS oligo(dT) primer and incubated at 72°C for 3 min, followed by reverse transcription using SMARTer II A oligo and SuperScript II reverse transcriptase for 90 min at 42°C and 10 min at 70°C. First-strand cDNA was purified using AMPure XP beads.

BCR heavy-chain sequences were amplified using the KAPA Real-Time Library Amplification Kit with 5PIIA forward and immunoglobulin reverse primers, with amplification monitored by real-time PCR and terminated during the exponential phase. Amplified products were purified using AMPure XP beads. Two subsequent PCR reactions were performed to add sample barcodes and Illumina sequencing adapters using KAPA HotStart ReadyMix, with AMPure XP purification following each amplification. Library quality was assessed using an Agilent Bioanalyzer, and libraries were sequenced on an Illumina MiSeq using paired-end sequencing. Primer sequences used for amplification and sequencing are provided in Supplementary Table X.

### BCR Sequence Analysis

Full-length V(D)J sequences were assembled de novo using Cell Ranger v7 for LN FNA samples and Cell Ranger v8 for PBMC samples. V, D, and J gene assignments were performed using IgBLAST with a database constructed from a previously described custom rhesus macaque germline V(D)J reference library^21^. IgBLAST output was parsed using the MakeDb.py igblast command in Change-O^63^ and converted to the Adaptive Immune Receptor Repertoire (AIRR) Community standard for downstream analysis within the Immcantation framework.

BM B_PC_ AIRR-seq reads were demultiplexed using a custom script and sequencing quality was assessed using FastQC v0.11.5. Reads were processed using the Immcantation pipeline v4.1.0. Paired-end reads were assembled using AssemblePairs with the --rc tail option and filtered using FilterSeq with a minimum quality score of 20. Forward primers were removed using MaskPrimers (--start 4, --mode cut), and duplicate sequences were collapsed using CollapseSeq (-n 20, --inner, --act set, --cf VPRIMER). Sequences were further filtered using SplitSeq (--num 2), annotated using IgBLAST v1.20.0 against the same custom rhesus macaque germline reference, and restricted to productive heavy-chain sequences using ParseDb.

Prior to combined repertoire analysis, BM B_PC_ sequences supported by two or fewer reads were excluded. Read counts within each B_PC_ sample were then normalized to the number of sorted cells, and sequences corresponding to less than one estimated cell equivalent after rounding were removed. This normalization was used only as a sequence-quality filter; subsequent B_PC_ analyses were performed using retained unique heavy-chain sequences without weighting by read abundance or estimated cell equivalents.

Only productive V(D)J sequences were retained for BCR repertoire analysis. Cells containing more than one productive heavy-chain contig were excluded from clonal analysis. B cell clones were defined from heavy-chain sequences using the scoper package. Pairwise distances between heavy-chain junction sequences were calculated using distToNearest with a length-normalized Hamming distance model, incorporating V- and J-gene assignments. The clonal distance threshold was determined from the nearest-neighbor distance distribution using findThreshold with the density method. Clones were subsequently assigned using hierarchicalClones at the resulting distance threshold, with clonal relationships restricted to sequences from the same animal.

Germline sequences were reconstructed using createGermlines with the custom rhesus macaque V(D)J reference library. Somatic hypermutation was quantified using SHazaM v1.1.2. For each cell, nucleotide mutations were counted across the aligned V(D)J region by comparing sequence_alignment with germline_alignment_d_mask using observedMutations (regionDefinition = NULL, frequency = FALSE). Replacement and silent mutations were quantified separately where indicated (combine = FALSE).

For phylogenetic analysis, clonal lineage trees were reconstructed using Dowser v2.1.0^64^. Maximum-likelihood trees were generated using getTrees (build = “pml”), and branch lengths were converted from substitutions per site to inferred mutation counts using scaleBranches (edge_type = “mutations”). The inferred germline sequence was included as the root of each lineage tree. Trees were visualized using the ape^65^ and ggtree^66^ packages.

### Clonal diversity analysis

Clonal diversity was assessed using the HC clonal assignments described above and was restricted to Env^+^ B_GC_ cells from LN FNA samples with an assigned clone. For each animal and time point, the number of cells belonging to each clone was determined. Clone number was defined as the total number of distinct clones recovered.

Clonal richness was estimated using the bias-corrected Chao1 estimator, which accounts for clones observed only once or twice to estimate the number of clones that may not have been sampled. Repertoire diversity was additionally assessed using the Gini–Simpson index (1 − D), calculated from the relative abundance of each clone using vegan v2.6-4.

Samples containing fewer than 30 Env^+^ B_GC_ sequences were excluded because of insufficient sampling depth. Two samples for which the standard deviation of the Chao1 estimate exceeded the estimate itself, reflecting a large excess of singleton over doubleton clones and correspondingly unstable extrapolation, were excluded from richness comparisons.

### Somatic hypermutation rate analysis

Analyses of SHM kinetics were restricted to Env⁺ B_GC_ cells from LN FNA samples with an assigned animal and mutation count. Mutation trajectories were anchored at zero mutations at week 0, reflecting the germline sequence at the time of immunization. Per-animal weekly mean mutation counts were calculated for animal–weeks containing at least three Env⁺ BGC cells, and a week-0 observation of zero mutations was included for each animal.

The breakpoint in mutation accumulation was identified by segmented regression (segmented v1.6-0) of pooled per-animal weekly means from Groups 2–4 over weeks 0–24, and departure from linearity was assessed using the Davies test. A segmented model fit comparably to smooth nonlinear alternatives (ΔAIC < 1 versus asymptotic and quadratic models) and substantially better than a linear model (ΔAIC = 25.6) and was selected because phase-specific mutation rates were the parameter of interest.

Phase-specific rates were then estimated for each group using a piecewise linear mixed-effects model (lme4 v1.1-35; lmerTest v3.1-3), with animal included as a random intercept and knots at weeks 9 and 24. The week-9 knot was based on the segmented regression, whereas the week-24 knot was imposed to reflect the booster schedule. Phase 1 versus Phase 2 was tested from the corresponding hinge coefficient, and Phase 1 versus Phase 3 from the linear combination of both hinge terms. Time points beyond week 48 were excluded because of limited sampling depth. Group differences were assessed using a single model containing group × phase interaction terms and no group main effect, thereby enforcing a shared week-0 intercept. The interaction was tested by likelihood-ratio comparison with a model containing common phase slopes, with both models refit by maximum likelihood.

Replacement and silent mutation rates were estimated per animal using the same piecewise framework. Because each animal contributed an independent slope without pooling across animals, these analyses were restricted to animal–weeks containing at least 20 Env⁺ B_GC_ cells. Rates were expressed as the percentage decrease from Phase 1 to Phase 2.

For clonal rate analyses, clones represented by at least five cells across at least two time points both before week 10 and between weeks 16 and 24 were selected. Phylogenetic trees were reconstructed as described above. Root-to-tip distances were calculated for each unique sequence using node.depth.edgelength in ape v5.7-1, with the inferred germline sequence assigned week 0 and a root-to-tip distance of zero. Per-clone mutation rates were estimated by piecewise regression through the origin, whereas clone-specific breakpoints were estimated by segmented regression with a free intercept. Because these distances were inferred from reconstructed phylogenies rather than counted directly from individual sequences, clonal and per-cell mutation rates were not numerically comparable.

The same analyses were applied to Env⁺ B_GC_ cells from a previously published ED-primed cohort^15^. The breakpoint was estimated independently for that cohort from its own segmented fit (week 7.9) rather than applying the breakpoint derived from the primary cohort.

Claude (Anthropic) was used to assist with implementation and debugging of R code for the SHM rate analyses. All statistical approaches, analysis code, and resulting outputs were reviewed and validated by the authors.

### Sequence mutability

Intrinsic sequence mutability was calculated for each cell using the S5F targeting model (calculateMutability, SHazaM) and expressed relative to the mutability of that cell’s own germline sequence. As no rhesus-specific targeting model is available, the human heavy-chain model (HH_S5F) was used^43^

### Single-cell Transcriptomic analysis

Gene expression, oligo-barcoded hashtag, and probe-count data were generated from the GEX and Feature Barcode libraries using Cell Ranger and aligned to the Ensembl Mmul10 reference genome, with mitochondrial genes incorporated from Mmul9. Gene expression data from sorted B cells isolated from both LN FNA and PBMC samples were initially analyzed together using Seurat v5^67^. Cells with fewer than 200 or more than 4,500 detected genes, or with >10% mitochondrial transcripts, were excluded. The proportions of mitochondrial, ribosomal, major histocompatibility complex (MHC), and immunoglobulin transcripts were additionally calculated for quality control. Rhesus macaque genes annotated with the *LOC* prefix, which included highly expressed immunoglobulin loci, were excluded from the RNA expression matrix before normalization to minimize the influence of immunoglobulin transcript abundance on normalization. Gene expression counts were log-normalized using NormalizeData, and variable features were identified using FindVariableFeatures. Feature Barcode counts, including oligo-barcoded sample hashtags and MD39 base-KO probe counts, were normalized using a centered log-ratio (CLR) transformation. Cells were then demultiplexed based on hashtag counts using MULTIseqDemux in Seurat v5. Samples were subsequently integrated within tissue using canonical correlation analysis (CCA) with FindIntegrationAnchors and IntegrateData to account for technical variation between sequencing runs and time points.

Following within-tissue integration, the FNA and PBMC datasets were merged into a combined Seurat object for initial quality control and assessment of B cell populations. The integrated expression assay was scaled and subjected to principal component analysis (PCA), followed by graph-based clustering and UMAP visualization using the first 15 principal components. Clustering was evaluated across resolutions from 0.1 to 1.0, with a resolution of 0.7 used for this initial analysis. V(D)J annotations generated independently using the IgBLAST/Change-O pipeline were then matched to the transcriptomic data by cell barcode. Clusters were inspected for expression of B cell and B_PC_ markers, transcriptomic quality, and productive V(D)J recovery. Clusters were excluded when they showed low expression of B cell markers, including *CD19* and *MS4A1*, in the absence of B_PC_ markers such as *XBP1* and *PRDM1*; contained disproportionately high mitochondrial or low ribosomal transcript abundance; or had poor V(D)J recovery, with >40% of cells lacking a productive V(D)J assignment. Cells classified as hashtag doublets or negatives were also excluded.

For the transcriptomic analyses presented in this study, LN FNA cells were then analyzed independently. RNA expression was log-normalized, and cell-cycle phase was assigned using CellCycleScoring with the Seurat S-phase and G2/M-phase gene sets. PCA, graph-based clustering, and UMAP visualization were performed using the integrated expression assay, with *LOC* and *MAMU* genes excluded from the features used for dimensionality reduction. Nearest-neighbor graphs and UMAPs were generated using the first 10 principal components. Clustering was evaluated across resolutions from 0.1 to 1.0, with a resolution of 0.4 used for the final FNA cluster assignments.

Clusters were annotated based on expression of canonical GC, memory B cell, and plasma cell marker genes together with gene set enrichment analysis (GSEA). Differentially expressed genes were identified using FindAllMarkers with a Wilcoxon rank-sum test, requiring expression in at least 25% of cells and an absolute log2 fold-change threshold of 0.25. GSEA was performed using the fgsea package with previously described human germinal center, memory B cell, and plasma cell transcriptional signatures ^68^. Where the published dataset contained multiple gene sets corresponding to the same cellular state, such as light zone (LZ), dark zone (DZ), memory B cell, or plasma cell populations, genes from the corresponding sets were combined into a single composite gene set for each state (**Table S3**). Genes were ranked by average log2 fold change for each cluster.

Based on transcriptional signatures and marker expression, 11 LN B cell populations were identified: five proliferating dark-zone populations (DZp1–5), a differentiating dark-zone population (DZd), three light-zone populations (LZ-Int, LZ, and LZ-Myc), an LN plasma cell population (LN B_PC_), and an LN memory B cell population (LN B_Mem_). Cells assigned to S or G2/M phase were classified as proliferating for subsequent analyses.

### Classification of base-binding and non-base-binding clones

Non-base-binding specificity was assessed using the oligo-barcoded MD39 base-KO probe. Sequencing-based probe counts were available for LN FNA samples at weeks 4, 8, 16, 27, 48, 55, and 59 and for pre- and post-boost PBMC samples; LN samples collected at weeks 10, 24, and 30 were not evaluable because only a fluorescent base-KO probe was used. Probe positivity was defined using a single threshold of 11 counts, corresponding to the 95th percentile of probe signal in week-4 Env⁻ cells (n = 8,781). Cells at or above this threshold were classified as non-base-binding.

Clone-level specificity was assigned by aggregating cell-level calls across all evaluable LN FNA and PBMC cells. To account for reduced probe detection in early low-affinity cells, positive calls were weighted by sampling time point (week 4, 0.5; week 8, 0.7; week 16, 0.8; all later time points and PBMC samples, 1.0). Clones with ≥3 evaluable cells were classified as non-base-binding when the weighted positivity score was ≥0.25 and as base-binding when <0.25. Clones represented by one or two evaluable cells were classified only when all calls were concordant; discordant clones and clones with no evaluable cells were left unclassified. Analyses comparing base- and non-base-binding clones were restricted to classified clones.

### ELISA

Plasma samples were thawed, heat-inactivated at 56°C for at least 30 min, and briefly centrifuged. Corning 96-well half-area plates were coated overnight at 4°C with streptavidin (2.5 μg/mL). Plates were washed three times with wash buffer (PBS containing 0.05% Tween-20) and coated with biotinylated MD39 WT (1 μg/mL) for 2 h at 37°C. Following three washes, plates were blocked with 3% BSA in PBS for 1 h at room temperature.

Plasma samples were serially diluted threefold in blocking buffer and incubated for 1.5 h at room temperature. Plates were then washed three times and incubated with goat anti-rhesus IgG-HRP (SouthernBiotech; 1:10,000 in blocking buffer) for 1 h at room temperature. After six washes, plates were developed with 1-Step Ultra TMB (Thermo Fisher Scientific), and the reaction was stopped with an equal volume of 2 N H₂SO₄ (Ricca Chemical). Absorbance at 450 nm was measured using an EnVision plate reader (PerkinElmer). Endpoint titers were interpolated in Prism 10 (GraphPad) using an asymmetric sigmoidal five-parameter logistic model of log-transformed concentration.

### Neutralization assays

Neutralization assays using the BG505.W6M.ENV.C2 isolate (T332N) pseudovirus were performed as previously described^16^. Assays were run blinded and in duplicate. Initial serum dilution was set at 1:10 and 50% inhibition (ID_50_) values below 30 were considered below the level of reliable measurement. Absolute ID50 values were calculated using normalized relative luminescence units and a customized nonlinear regression model with the bottom constraint set to 0 and top constraint set to the <100.

### ELISpot

Bone marrow antibody-secreting cells (ASCs) were quantified by enzyme-linked immunosorbent spot (ELISpot) assay as previously described^36^. Briefly, 96-well MultiScreen HTS filter plates (MSHAN4B50, Millipore) were coated overnight at 4°C with either goat anti-human Ig (5 µg/mL; SouthernBiotech, cat. no. 2010-01) to quantify total rhesus IgG-secreting ASCs or Galanthus nivalis lectin (GNL; 20 µg/mL; Vector Laboratories, cat. no. L-1240) for capture of MD39 SOSIP Env. Following washing, plates were blocked with R10 medium for 2 h at 37°C. GNL-coated wells were subsequently incubated with MD39 SOSIP Env (10 µg/mL) for 2 h at 37°C.

BM mononuclear cells were resuspended at 5 × 10⁶ cells/mL, plated in duplicate, serially diluted threefold, and incubated for 16 h at 37°C in 5% CO₂. Following washing with PBS and PBS containing 0.05% Tween-20, captured IgG was detected using biotin-conjugated goat anti-human IgG (1:500; SouthernBiotech, cat. no. 2045-08), followed by Avidin D-HRP (1:1,000; Vector Laboratories). Spots were developed using 3-amino-9-ethylcarbazole (AEC) substrate and imaged using an ImmunoSpot CTL analyzer with Image Acquisition 4.5 software (Cellular Technology). Total IgG⁺ and MD39 SOSIP-specific ASCs were quantified as spots per 10⁶ BM mononuclear cells, and antigen-specific responses were expressed as the percentage of MD39 SOSIP-specific ASCs among total IgG⁺ ASCs.

### EMPEM

Plasma antibodies were purified using Protein A Sepharose resin (GE Healthcare), eluted from the resin with 0.1 M glycine at pH 2.5, and buffer exchanged into 1× PBS. Fabs were generated using crystalline papain (Thermo Scientific), digested for 5 h at 37 °C and purified using Protein A Sepharose resin to remove Fc and undigested IgG. The flow through containing polyclonal Fabs was buffer exchanged into 1× Tris-buffered saline (TBS). Complexes were assembled with 1.0 mg of polyclonal Fabs incubated overnight with 15 µg of MD39 Env trimers at room temperature, followed by purification to remove unbound Fab by size-exclusion chromatography (SEC) using a Superose 6 Increase 10/300 column (GE Healthcare). Complexes were diluted to 20 µg ml^−1^, immediately placed on 400-mesh Cu grids and stained with 2% (w/v) uranyl formate for 40 s. Imaging was performed using a Thermo Fisher Scientific Glacios transmission electron microscope operating at 200 kV, equipped with Thermo Fisher Scientific Falcon 4 camera (73,000× magnification, 1.89 Å pixel size). Automated data collection was performed using EPU (Thermo Fisher Scientific) and data processing was performed using RELION 4.0^69^, following standard procedures for reference-free particle picking and 2D classification. Following three rounds of 2D classification, particles from classes corresponding to immune complexes were subjected to 3D refinement with C3 symmetry and a 40 Å low-pass filtered map of HIV Env ectodomain as the initial model. The initial model is based on PDB coordinates 6V0R, converted <u>to</u> a map using the molmap feature in UCSF ChimeraX^70^. Following 3D refinement, C3 symmetry expansion was applied to the particles and 8 separate focused 3D classification “skip align” jobs were run (K=10), each with a 40 Å diameter spherical mask over key HIV Env epitopes.

The names of the epitopes and reference structures used for orienting the masks are: 1) gp41-base (PDB 7L8Z), 2) gp41-GH (PDB 7L8U), 3) gp41-FP (PDB 7L8T), 4) gp120-GH (PDB 7L8B), 5) C3/V5 (PDB 7L8Y), 6) CD4bs and gp120 interface (PDB 7L8X), 7) V1/V3 (PDB 7L8E) and 8) V2-apex (PDB 9D1W). For each epitope 3D classification, classes with visible Fab density were selected and subjected to 3D refinement, 2D classification, and a second round of 3D classification. If the 3D refinement resulted in partial Fab density relative to the Env trimer, classes were selected from the subsequent round of 3D classification, and this was repeated until the reconstruction improved or no change was noted. Final reconstructions were visually inspected and assigned the correct epitope label. The number of final particles belonging to each epitope was divided by the total number of particles in the initial (C3) 3D refinement. This value, which can range from 0 to 3 due to C3 symmetry expanded particles used in the 3D classification steps, is described as the *EMPEM magnitude.* The *Total Magnitude* is the sum of each individual epitope *EMPEM magnitude* for a given animal and time point. Representative maps have been deposited to the Electron Microscopy Data Bank. UCSF ChimeraX was used to visualize and segment 3D-refined maps.

### Statistical analysis

All statistical analyses were performed using GraphPad Prism 10 (GraphPad Software) or R (v4.3.2). Statistical tests, multiple-comparisons corrections, and summary statistics for individual analyses are indicated in the corresponding figure legends. Specialized analyses of somatic hypermutation kinetics, including segmented regression and linear mixed-effects modeling, are described separately above. Data are presented as medians, mean ± SEM, or model estimates with 95% confidence intervals, as indicated in the figure legends. Statistical significance is indicated as follows: *P* < 0.05 (\**), P < 0.01 (**), P < 0.001 (\*\*\**), and *P* < 0.0001 (****); non-significant comparisons are not shown unless otherwise indicated.

## SUPPLEMENTARY TABLES

**Supplementary Table 1: Bilateral sharing of Env-specific germinal center B cell clones between the left and right lymph node**

**Supplementary Table 2: Flow cytometry and FACS antibody panels and reagents**

**Supplementary Table 3: Gene sets used for GSEA**

## SUPPLEMENTARY FIGURE LEGENDS

**Supplementary Figure 1.**
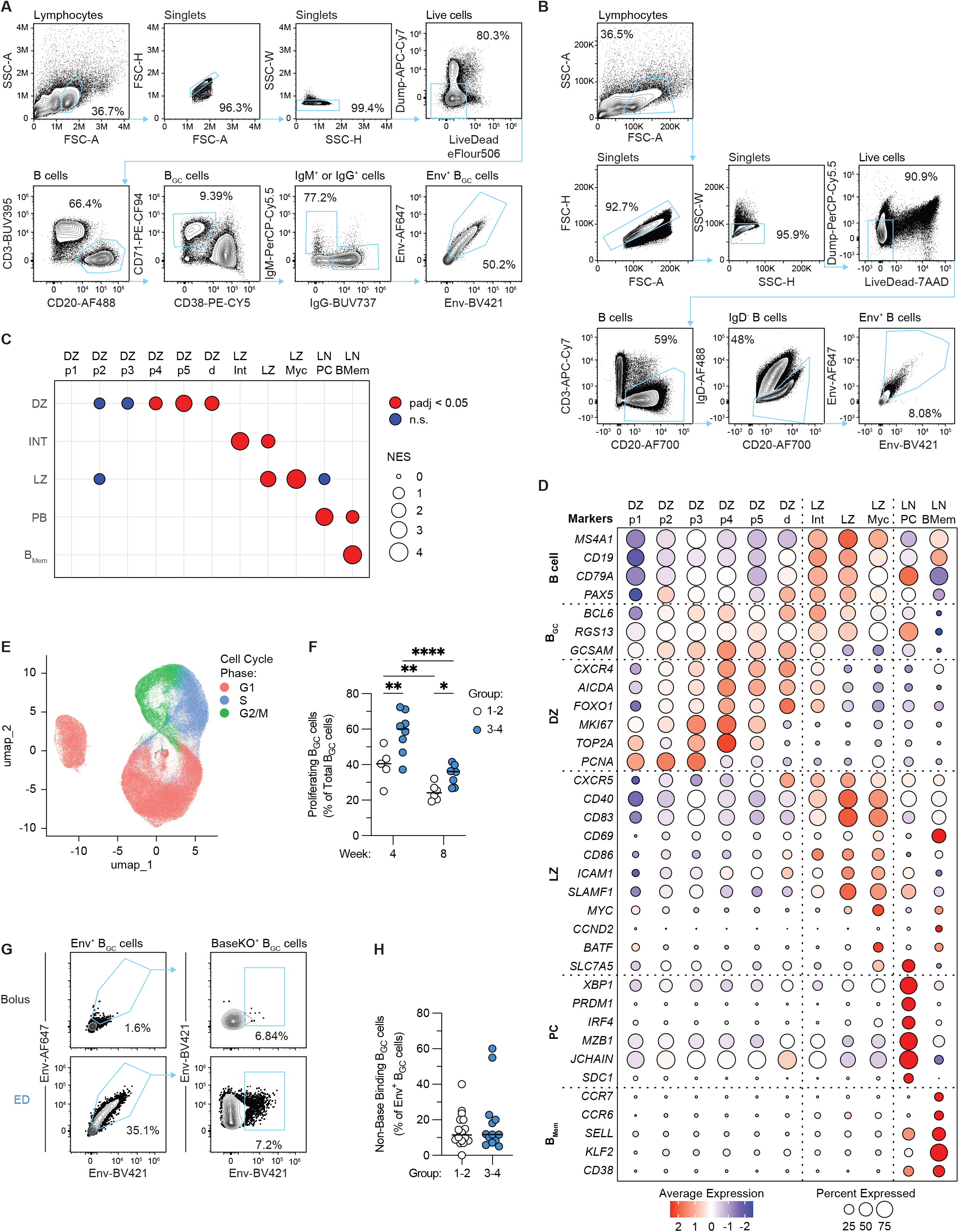
Flow cytometry gating and transcriptional characterization of LN B cell populations. **(A)** Representative flow cytometry gating strategy used to identify Env⁺ B_GC_ cells in LN FNAs. **(B)** Representative flow cytometry gating strategy used to sort CD20⁺IgD⁻ Env⁺ B cells for single-cell RNA and paired VDJ sequencing for LN FNAs. **(C)** Gene set enrichment analysis (GSEA) of the 11 LN B cell clusters identified by single-cell RNA sequencing. Gene sets corresponding to dark zone (DZ), intermediate (Int), light zone (LZ), plasma cell (B_PC_), and memory B cell (B_Mem_) states were derived from Holmes et al 2020. Where multiple gene sets represented the same cellular state, genes were combined into a single gene set before analysis. Dot size indicates the normalized enrichment score (NES); red indicates statistically significant enrichment and blue indicates non-significant enrichment. **(D)** Relative expression of selected B cell, B_GC_, DZ, LZ, B_PC_, and B_Mem_ marker genes across the 11 LN B cell clusters. Dot size indicates the percentage of cells expressing each gene, and color indicates relative expression. **(E)** UMAP of LN B cells shown in Fig. 1D, colored by inferred cell-cycle phase. **(F)** Frequency of proliferating B_GC_ cells, defined as cells in S or G2/M phase, as a percentage of total B_GC_ cells at weeks 4 and 8 following bolus (Groups 1 and 2) or ED (Groups 3 and 4) priming. Samples with fewer than 30 Env⁺ B_GC_ sequences were excluded because of insufficient sampling depth. **(G)** Representative flow cytometry gating strategy using the BaseKO probe to identify non-base-binding cells among Env⁺ B_GC_ cells at week 10. **(H)** Frequency of non-base-binding B_GC_ cells among Env⁺ B_GC_ cells at week 10 following bolus or ED priming. Statistical comparisons were performed using two-way ANOVA with multiple-comparisons correction for **(F)** and a two-tailed Mann–Whitney test for **(H)**. *P* values are indicated as *P* < 0.05 (\**), P < 0.01 (**), P < 0.001 (\*\*\**), and *P* < 0.0001 (****); non-significant comparisons are not shown. Horizontal bars indicate median.

**Supplementary Figure 2.**
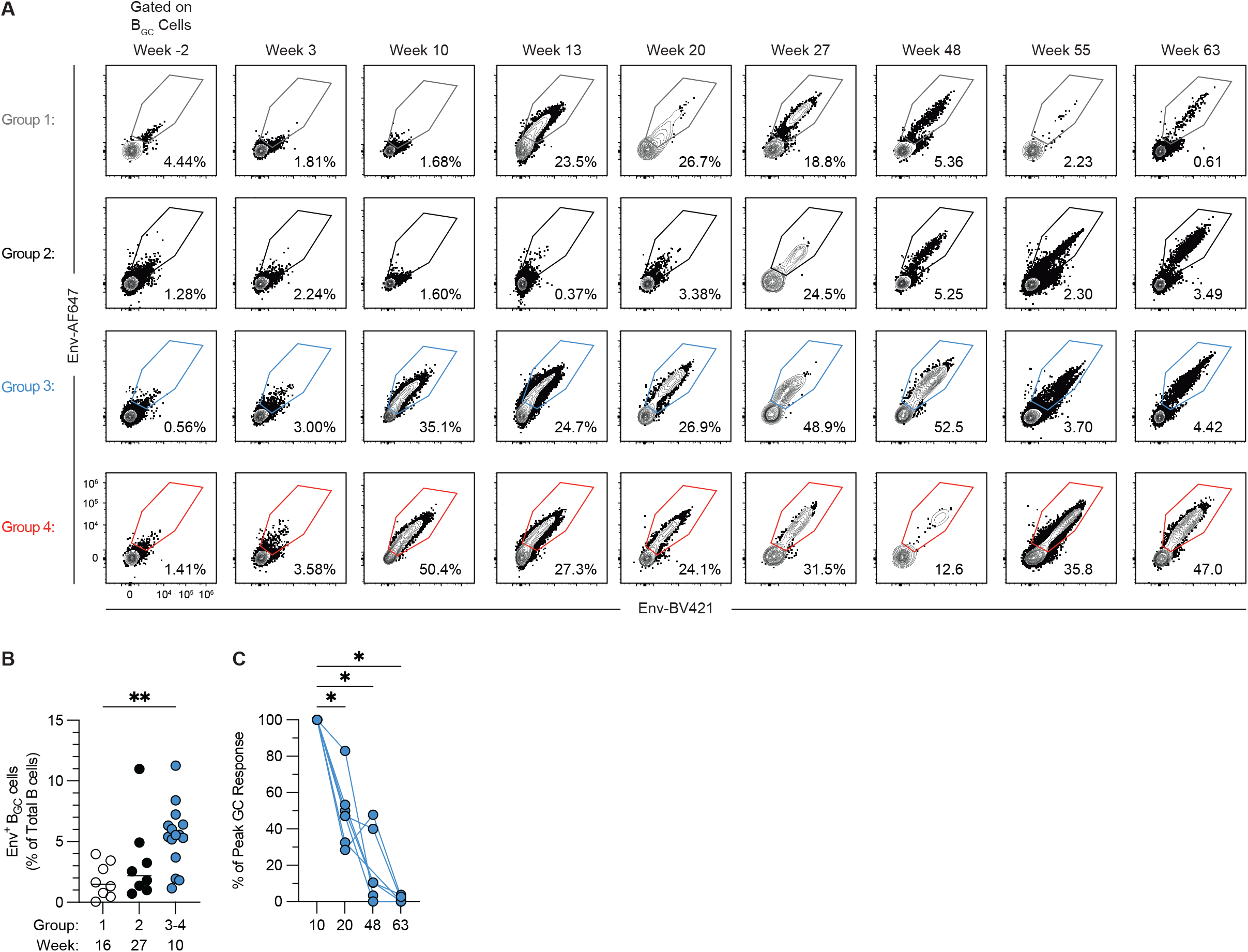
Longitudinal persistence of Env⁺ GC responses. **(A)** Representative flow cytometry plots showing Env⁺ B_GC_ cells from one animal per group at each time point shown in Fig. 2A. **(B)** Frequency of Env⁺ B_GC_ cells as a percentage of total B cells at the peak GC response for each immunization regimen: week 16 for Group 1, week 27 for Group 2, and week 10 for the combined ED-primed Groups 3 and 4. **(C)** Persistence of the Env⁺ B_GC_ cell response in Group 3 animals at weeks 20, 48, and 63, expressed relative to each animal’s peak week-10 response after baseline correction (baseline = 0%; week 10 = 100%). Statistical comparisons were performed using a Kruskal–Wallis test with multiple-comparisons correction for **(B)** and two-tailed Wilcoxon matched-pairs signed-rank tests for **(C)**. P values are indicated as P < 0.05 (*), P < 0.01 (**), P < 0.001 (***), and P < 0.0001 (****); non-significant comparisons are not shown. Horizontal bars indicate median.

**Supplementary Figure 3.**
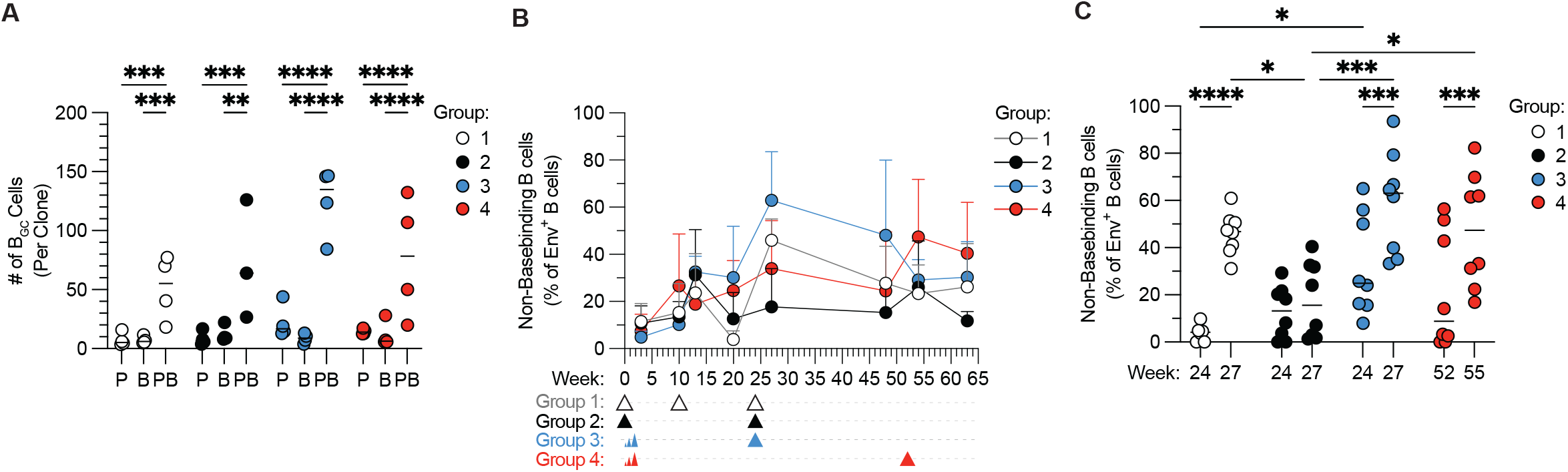
Clonal and epitope-specificity dynamics of prolonged GC responses. **(A)** Average number of Env⁺ B_GC_ cells per clone among clones detected during priming only (Prime; P), following boosting only (Boost; B), or during both periods (Prime + Boost; PB). **(B)** Frequency of non-base-binding B_GC_ cells among Env⁺ B_GC_ cells over time. **(C)** Frequency of non-base-binding B_GC_ cells among Env⁺ B_GC_ cells before and after boosting. Statistical comparisons were performed using two-way ANOVA with multiple-comparisons correction for **(A)** and **(C)**. For **(A)**, statistical comparisons were performed between clone categories within each group; no comparisons were performed between groups. P values are indicated as P < 0.05 (*), P < 0.01 (**), P < 0.001 (***), and P < 0.0001 (****); non-significant comparisons are not shown. Horizontal bars indicate median.

**Supplementary Figure 4.**
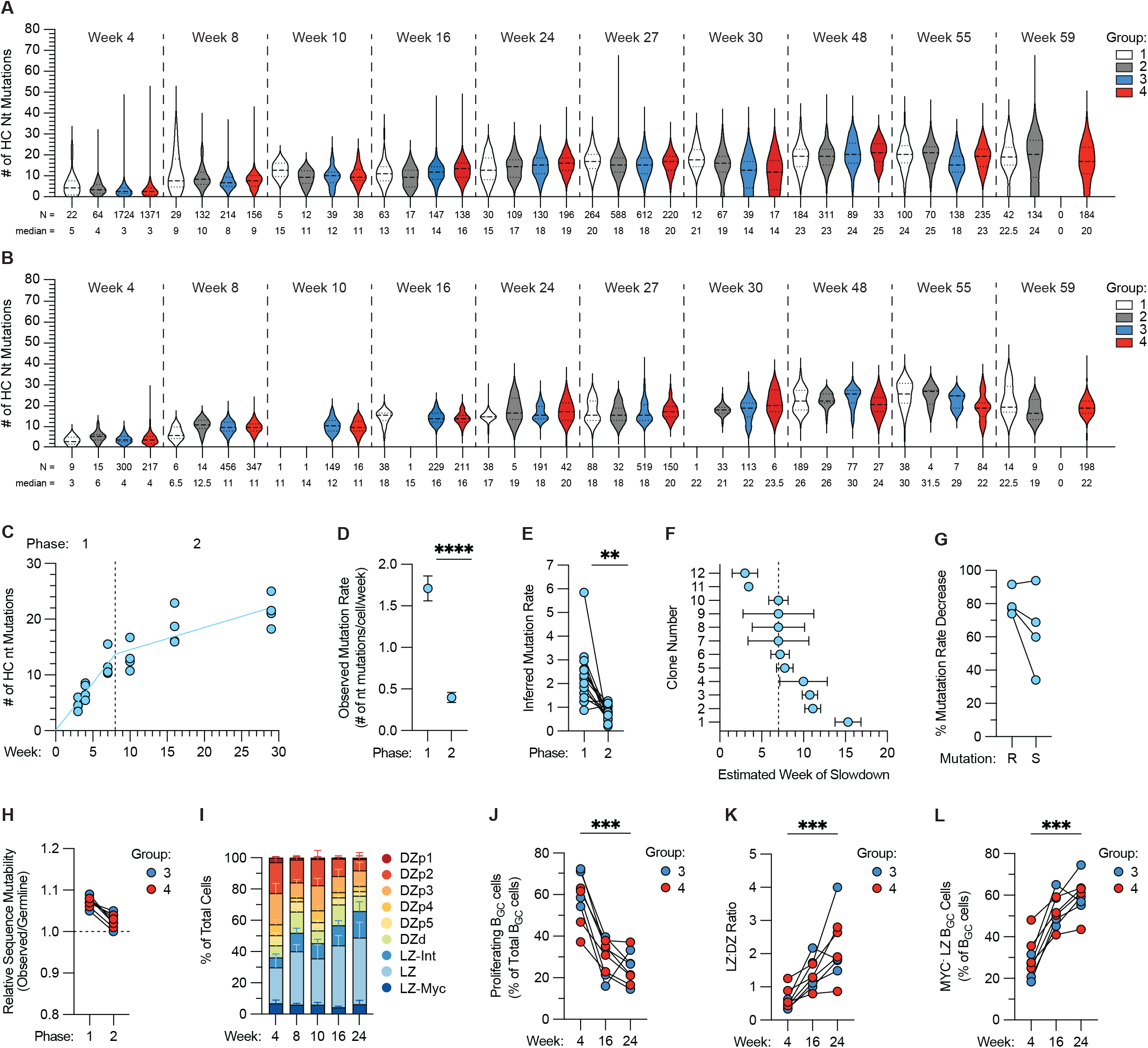
Somatic hypermutation kinetics and cellular features during prolonged GC responses. **(A)** Heavy-chain (HC) nucleotide (Nt) mutations per Env⁺ LN B_Mem_ cell over time. Horizontal lines indicate the median and quartiles. The number of cells (N) and median mutation count are shown below each violin. **(B)** HC Nt mutations per Env⁺ LN B_PC_ over time. Horizontal lines indicate the median and quartiles. The number of cells (N) and median mutation count are shown below each violin. **(C)** Mean HC Nt mutations per animal over time in an independent previously published cohort. Lines show the fitted two-phase trajectory from a piecewise linear mixed-effects model fit to per-animal weekly means, with each animal– week contributing equally. The vertical dotted line marks the week 8 breakpoint between phase 1 (weeks 0–8) and phase 2 (weeks 8–29), as determined by segmented regression of the pooled data. **(D)** Observed mutation rate (HC Nt mutations per cell per week) during phases 1 and 2 in the independent cohort, derived from the model in (C). Error bars indicate 95% confidence intervals. **(E)** Inferred mutation rate per clone during phases 1 and 2 in the independent cohort; lines connect phases within a clone. Per-clone rates were obtained by piecewise regression of inferred mutations from germline against sampling week, with the breakpoint set at week 8. **(F)** Estimated breakpoint (week of slowdown) for the 12 persistent clones identified in the independent cohort. The dotted line indicates the median week of slowdown (7). Error bars indicate 95% confidence intervals. **(G)** Percentage decrease in observed mutation rate from phase 1 to phase 2 for replacement (R) and silent (S) mutations per animal in the independent cohort. **(H)** Relative sequence mutability of Env⁺ B_GC_ cells in ED-primed animals (Groups 3 and 4) during phase 1 (weeks 0–9) and phase 2 (weeks 9–24). Lines connect phases within an animal. Values are expressed relative to each cell’s germline sequence; the dashed line at 1.0 indicates mutability unchanged from germline. **(I)** Relative abundance of LN B_GC_ cell clusters in Groups 3 and 4 over time. **(J)** Frequency of proliferating B_GC_ cells, defined as cells in S or G2/M phase, as a percentage of total B_GC_ cells at weeks 4, 16, and 24 following ED priming (Groups 3 and 4). **(K)** Light zone-to-dark zone (LZ:DZ) ratio of Env⁺ B_GC_ cells over time following ED priming. **(L)** Frequency of the MYC⁻ LZ population (LZ and LZ-Int clusters) among B_GC_ cells over time following ED priming. Statistical comparisons were performed using Wilcoxon matched-pairs signed-rank tests for **(D, E and G)** and Friedman tests with multiple-comparisons correction for **(J-L)**. P values are indicated as P < 0.05 (*), P < 0.01 (**), P < 0.001 (***), and P < 0.0001 (****); non-significant comparisons are not shown. Horizontal bars indicate median.

**Supplementary Figure 5.**
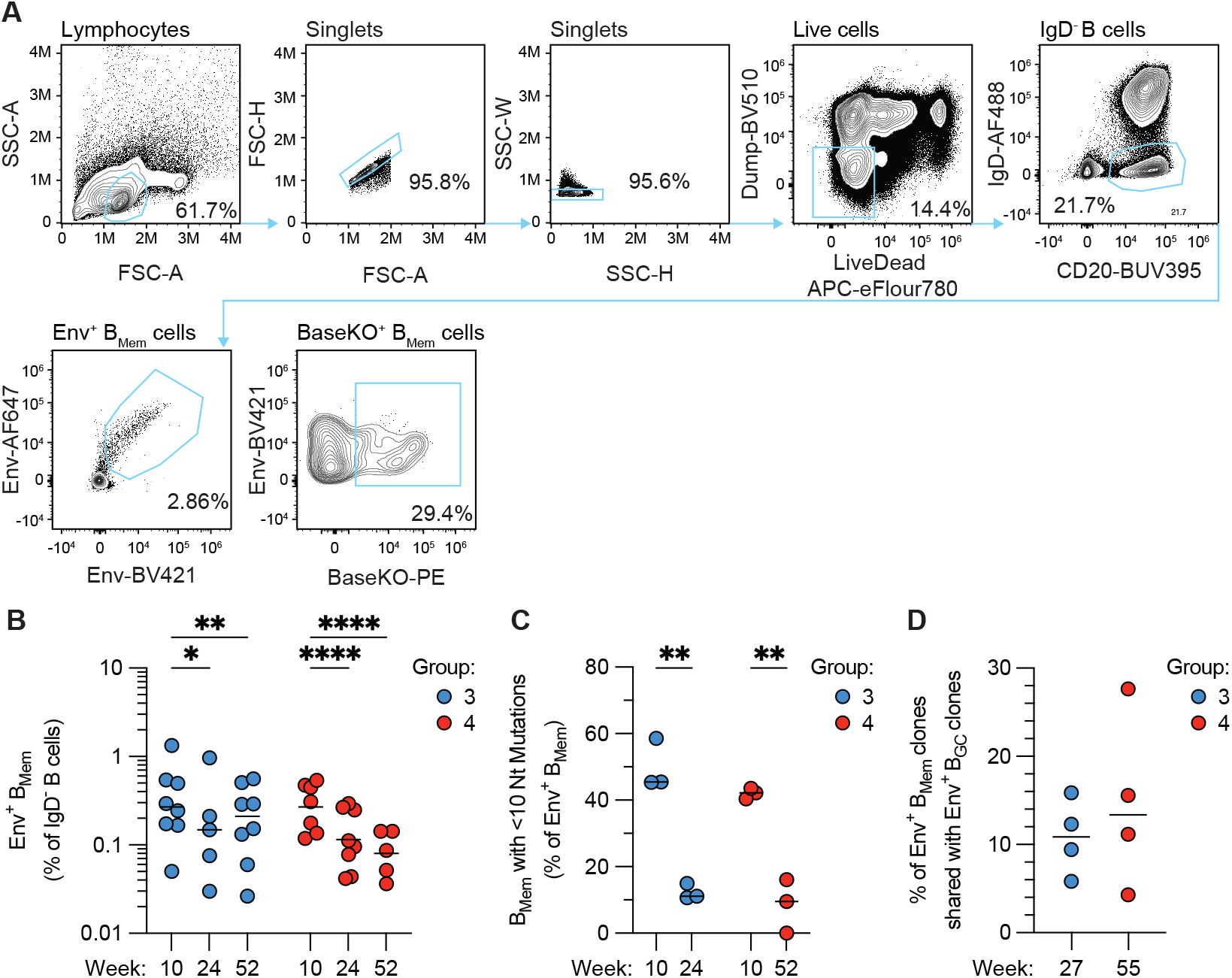
Dynamics and clonal relationships of Env⁺ B_Mem_ cells during prolonged GC responses. **(A)** Representative flow cytometry plots showing the gating strategy used to identify and sort Env⁺ B_Mem_ cells in peripheral blood. **(B)** Frequency of Env⁺ B_Mem_ cells among total IgD⁻ B cells over time in ED-primed animals (Groups 3 and 4). **(C)** Frequency of Env⁺ B_Mem_ cells with ≤10 HC Nt mutations at weeks 10 and 24 in ED-primed animals (Groups 3 and 4). One animal from each group was excluded because no Env⁺ B_Mem_ sequences were recovered at either time point. **(D)** Percentage of post-boost Env⁺ B_Mem_ clones that shared a clonal lineage with Env⁺ B_GC_ clones present before boosting. Statistical comparisons were performed using two-way ANOVA with multiple-comparisons correction for **(B, C)** and a Mann–Whitney test for **(D)**. P < 0.05 (*), P < 0.01 (**), P < 0.001 (***), P < 0.0001 (****); non-significant comparisons are not shown. Horizontal bars indicate median.

**Supplementary Figure 6.**
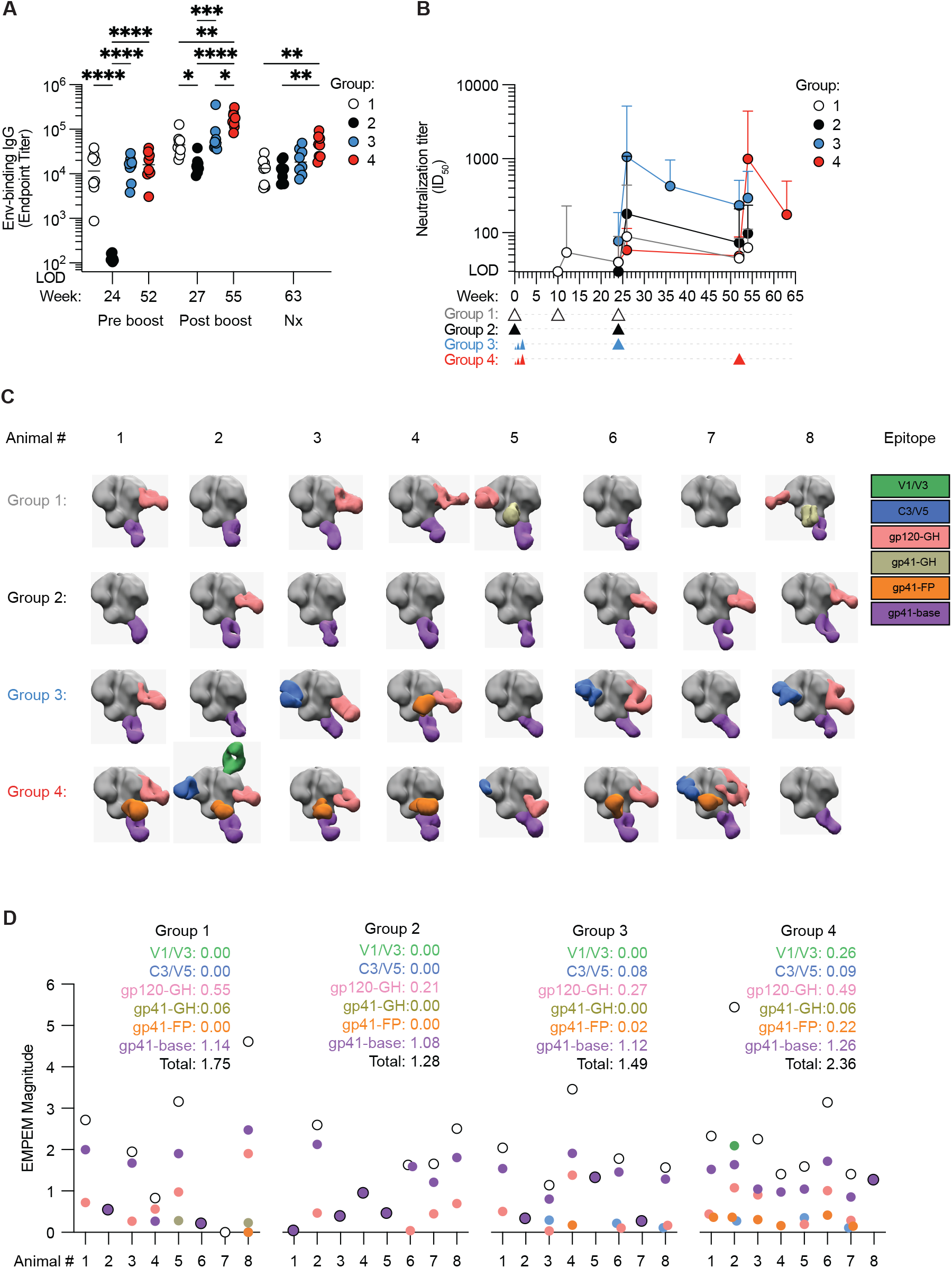
Longitudinal serum antibody responses and EMPEM analysis following boosting. **(A)** Env-specific serum IgG endpoint titers before boosting (week 24 for Groups 1–3; week 52 for Group 4), two weeks after boosting (week 27 for Groups 1–3; week 55 for Group 4), and at necropsy (week 63 for all groups). For Group 1, pre- and post-boost measurements correspond to the second boost. The y-axis begins at the limit of detection (LOD). **(B)** Autologous neutralizing antibody titers (ID₅₀) over time, colored by group. The y-axis begins at the LOD. **(C)** EMPEM reconstructions showing Env trimers in gray and bound antibody Fabs colored according to the Env epitope recognized. **(D)** EMPEM magnitude two weeks after boosting, representing the average number of antibody Fabs bound per Env trimer. Magnitudes are shown for each epitope and for the total response in each animal. Filled circles indicate individual epitope-specific magnitudes, and open circles indicate the total magnitude summed across all detected epitopes. When only a single epitope was detected, the total and epitope-specific values are identical and the symbols are superimposed, appearing as a colored circle with a black outline. Mean EMPEM magnitude for each study group is listed by epitope and for the total response. Colors correspond to the epitope labels shown in (C). Statistical comparisons were performed using two-way ANOVA with multiple-comparisons correction for **(A)**. P values are indicated as P < 0.05 (*), P < 0.01 (**), P < 0.001 (***), and P < 0.0001 (****); non-significant comparisons are not shown. Horizontal bars indicate median.

**Supplementary Figure 7.**
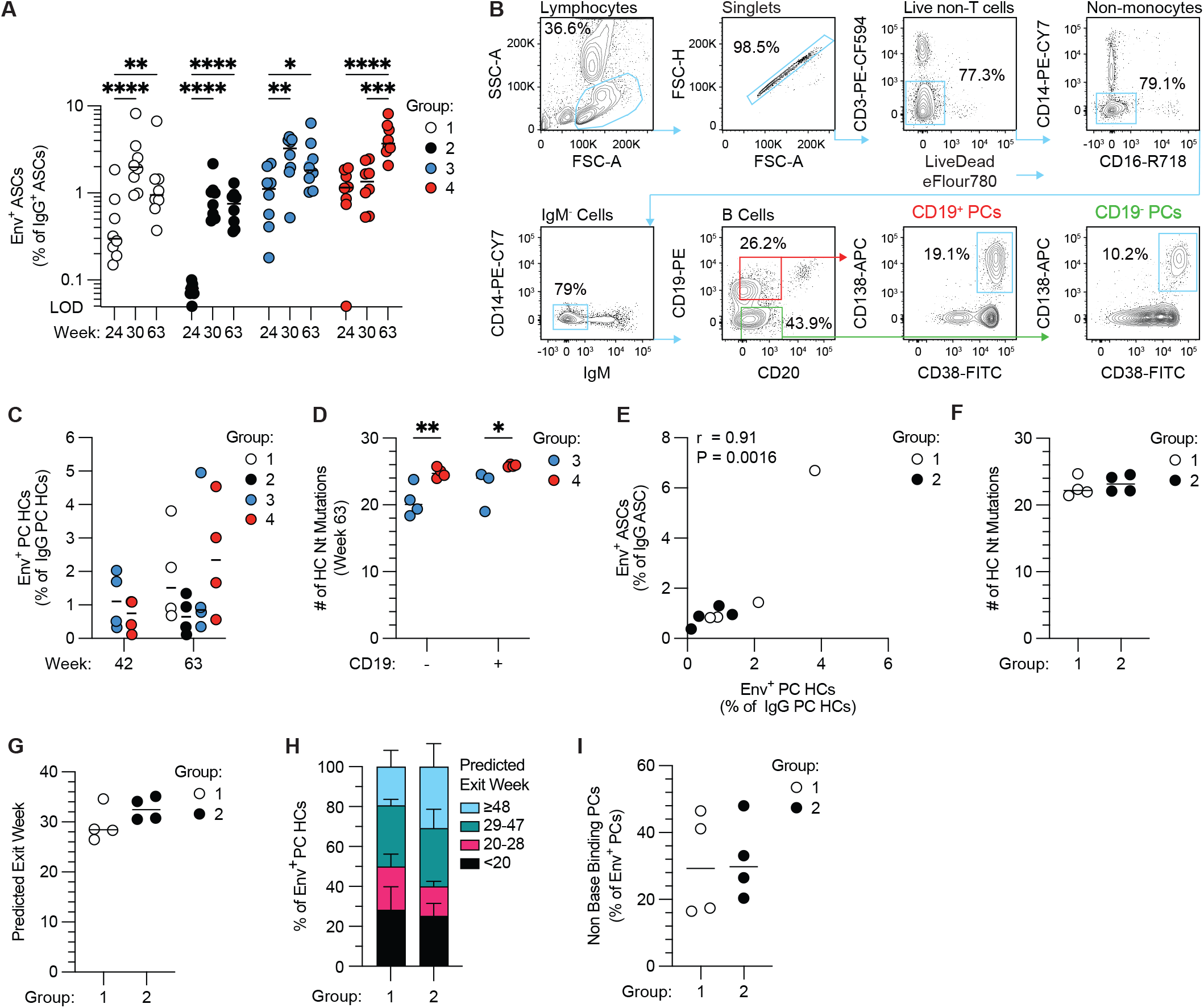
BM B_PC_ responses and inferred timing of B_PC_ generation. **(A)** Env⁺ ASCs as a percentage of total IgG⁺ ASCs in BM at weeks 24, 30, and 63. The y-axis begins at the limit of detection (LOD). **(B)** Representative flow cytometry gating strategy used to identify and sort BM B_PCs_. **(C)** Mean number of HC Nt mutations per Env⁺ BM B_PC_ sequence in CD19⁺ and CD19⁻ B_PCs_ from ED-primed Groups 3 and 4 at week 63. **(D)** Number of Env⁺ BM B_PC_ HC sequences identified by clonal overlap with Env⁺ B_GC_ or B_Mem_ cells at week 42 in ED-primed animals (Groups 3 and 4) and at week 63 in all groups. **(E)** Correlation between Env⁺ ASCs as a percentage of total IgG⁺ ASCs, determined by ELISpot, and Env⁺ B_PC_ HC sequences identified by clonal overlap with Env⁺ B_GC_ or B_Mem_ cells, as a percentage of total IgG B_PC_ HC sequences, shown for bolus-primed Groups 1 and 2. *r* indicates the Spearman correlation coefficient. **(F)** Mean number of HC Nt mutations per Env⁺ BM B_PC_ sequence for each animal in bolus-primed Groups 1 and 2 at week 63. **(G)** Mean predicted GC exit week of Env⁺ BM B_PC_ sequences for each animal in bolus-primed Groups 1 and 2 at week 63. **(H)** Distribution of Env⁺ BM B_PC_ HC sequences predicted to have exited the GC before week 20, between weeks 20– 28, between weeks 29–47, or at or after week 48, shown for bolus-primed Groups 1 and 2 at week 63. Bars indicate mean ± SEM. **(I)** Frequency of non-base-binding B_PCs_ among Env⁺ BM B_PCs_ at week 63 in bolus-primed Groups 1 and 2. Statistical comparisons were performed using two-way ANOVA with multiple-comparisons correction for **(A),** Spearman correlation for **(E)**, and two-tailed Mann–Whitney tests for **(F–I)**. P values are indicated as P < 0.05 (*), P < 0.01 (**), P < 0.001 (***), and P < 0.0001 (****); non-significant comparisons are not shown. Horizontal bars indicate median.

